# Caspase-11 Mediated Hyperinflammation Impairs CD8⁺ T Cell Immunity and Viral Clearance in Severe SARS-CoV-2 Infection

**DOI:** 10.1101/2025.11.01.684729

**Authors:** Mostafa M. Eltobgy, Mohamed M. Shamseldin, Owen Whitham, Heba M. Amer, Jeffery Atkinson, Asmaa Badr, Jesse Hall, Jihad Omran, Gauruv Gupta, Yara Hassan, Rabab Abdelaleem, Richard Perez, Sarah Faber, Maciej Pietrzak, Amy Webb, Xiaoli Zhang, Adam D. Kenney, Destiny Bissel, Shady Estfanous, Kylene Daily, Andrew Mcnamara, KC Mahesh, Mark E. Peeples, Emily A. Hemann, Shahid M. Nimjee, Estelle Cormet-Boyaka, Jianrong Li, Prosper N. Boyaka, Jacob S. Yount, Benjamin Segal, Purnima Dubey, Amal O. Amer

## Abstract

Severe SARS-CoV-2 infection is characterized by lung hyperinflammation, impaired interferon responses, and defective T-cell activation, yet the molecular drivers of these immune dysregulations remain incompletely understood. Caspase-11 (CASP11), a key mediator of the non-canonical inflammasome, has been shown to mediate an innate hyperinflammatory response and cytokine release in a non-severe, non-lethal SARS-CoV-2 infection model. However, the role played by CASP11 in severe SARS-CoV-2 disease and how it impacts adaptive immunity is not identified. Here, we newly discover that CASP11 exacerbates severe SARS-CoV-2 pathogenesis by amplifying early innate immune responses while concurrently impairing antiviral CD8 T-cell immunity. Using global knockouts, reciprocal bone marrow chimeras, and phagocyte-monocyte system (PMS) cell-specific CASP11 deletion models, we show that CASP11 deletion in monocyte-derived cells reduces lung inflammation, enhances type I and II interferon signaling, and promotes robust virus-specific effector CD8⁺ T-cell response. This was associated with enhanced viral clearance and improved survival, even under lethal infection conditions. Importantly, CASP11 KO mice also exhibited faster resolution of post-viral inflammation, suggesting a role in long-term immune remodeling. These findings position CASP11 as a promising immunomodulatory target for acute and delayed manifestations of severe SARS-CoV-2.

## Introduction

SARSl1CoVl12, the etiologic agent of Covid-19, is an enveloped, positivel1sense, singlel1stranded RNA virus belonging to the betacoronavirus genus(1). Although most SARS-CoV-2 infections result in mild, selfl1limited upper respiratory symptoms, a significant subset of individuals develop severe respiratory disease that can rapidly progress to acute respiratory distress syndrome (ARDS), respiratory failure, and multi-organ damage. In unvaccinated individuals, this infection has caused millions of deaths worldwide(2–4).

Severe SARS-CoV-2 disease is associated with an initial exaggerated innate immune response characterized by excessive release of pro-inflammatory cytokines such as IL-1β, IL-6, and IL-8, often referred to as a "cytokine storm," accompanied by significant neutrophil infiltration into the lungs(5,6). This inflammatory response mediates lung tissue damage and can result in respiratory failure and systemic inflammation(7). However, there are other cytokines that have been shown to be suppressed in the setting of severe SARS-CoV-2 infection. Specifically, type I interferon activity is diminished in critically sick SARS-CoV-2 patients, which was associated with persistent viremia and an exacerbated inflammatory response(8). Additionally, interferon-gamma responses are compromised in these patients(9,10). This inadequate interferon response, along with the plethora of proinflammatory cytokines, interferes with initiating an adequate and effective adaptive immune response in patients with severe SARS-CoV-2 disease(2,11).

Patients with severe SARS-CoV-2 infection exhibit suboptimal T cell responses, marked by quantitative and qualitative alterations in both CD4⁺ and CD8⁺ T cell compartments(12–19). Several clinical studies have reported significant reductions in overall lymphocyte counts (lymphopenia), which correlates with disease severity and poor clinical outcomes(20,21). Additionally, severe SARS-CoV-2 infection is associated with notable phenotypic alterations within the T cell pool, including increased T cell exhaustion, impaired activation and cytokine production, and disrupted T cell subset distribution(17,18). However, the exact mechanisms driving T cell response impairment remain unclear. These alterations could result from direct viral invasion, prolonged antigen exposure, or indirectly through virus-induced excessive innate immune activation. Limited mechanistic and longitudinal studies have hindered a comprehensive understanding of these processes.

The inflammasome is a critical component of the innate immune system which plays a major role in SARS-CoV-2 infection(22–27). Among the best-characterized inflammasomes is the NLRP3 inflammasome, which is activated by a range of viral and host-derived danger signals during SARS-CoV-2 infection. Upon activation, NLRP3 recruits the adaptor protein ASC, leading to the activation of Caspase-1, a canonical inflammasome effector protease. Caspase-1 cleaves pro-inflammatory cytokines IL-1β and IL-18 into their active forms and induces pyroptosis, a form of inflammatory cell death, thereby shaping the early inflammatory phase of SARS-CoV-2 infection(26,28,29).

Caspase-11(CASP11), and its human homologs caspasel14/5, is a critical member of the nonl1canonical inflammasome(30,31). We previously showed that the lack of CASP11 decreases the innate inflammatory responses during moderate SARS-CoV-2 infection without affecting viral replication in the lung(32). In contrast, the lack of other inflammasome components such as NLRP3 or caspasel11 attenuate inflammation but promotes viral replication(27). CASP11 activation significantly amplifies inflammation by inducing cytokine production and pyroptotic cell death, suggesting its potential role in dysregulating subsequent adaptive immune responses(32–34). We previously demonstrated that CASP11 mediates cytokine release, neutrophil infiltration, and neutrophil extracellular trap (NET) formation after moderate SARS-CoV-2 infection(32). Additionally, we observed elevated CASP11 expression in lung tissue from humans with severe SARS-CoV-2 infection(32). However, the role of CASP11 in severe SARS-CoV-2 infection has not been well investigated, particularly with regard to adaptive immune responses. Here, we find that a CASP11-mediated innate immuno-inflammatory response compromises an effective adaptive T-cell response, contributing to severe disease outcomes in SARS-CoV-2 infection.

We mechanistically define that CASP11 mediates its effects on both innate and adaptive immune responses after SARS-CoV-2 infection through its role in monocyte-derived cells. It exacerbates lung inflammation, suppresses type I and II interferon signaling, and hinders the development of robust virus-specific effector CD8⁺ T cell responses.

## Results

### CASP11 deficiency protects against severe SARS⍰CoV⍰2 infection, limits early hyperinflammation and enhances interferon responses

We infected 12-16l1weekl1old WT and CASP11 KO (*Casp11*⁻^/^⁻) mice with 5*10^5 TCID50 of the MA10 mousel1adapted SARS-CoV-2 virus(35). WT mice developed severe disease, losing approximately 30% of their body weight by day 7 post-infection with no signs of recovery ***(Fig. 1a).*** In contrast, *Casp11*⁻^/^⁻ mice showed significantly milder disease, with recovery and notable weight regain starting at day 4 and continuing through day 7 post-infection ***(Fig. 1a).*** We next evaluated pulmonary function using whole-body plethysmography, a non-invasive method that measures PenH as a surrogate indicator of airway resistance in conscious, unrestrained mice. *Casp11*⁻^/^⁻ mice exhibited significantly lower PenH compared to WT, indicating less respiratory distress and better lung function during infection **(Fig. 1b).** This shows that in the absence of CASP11, mice are protected against clinically severe SARS-CoV-2 infection.

**Figure 1.**
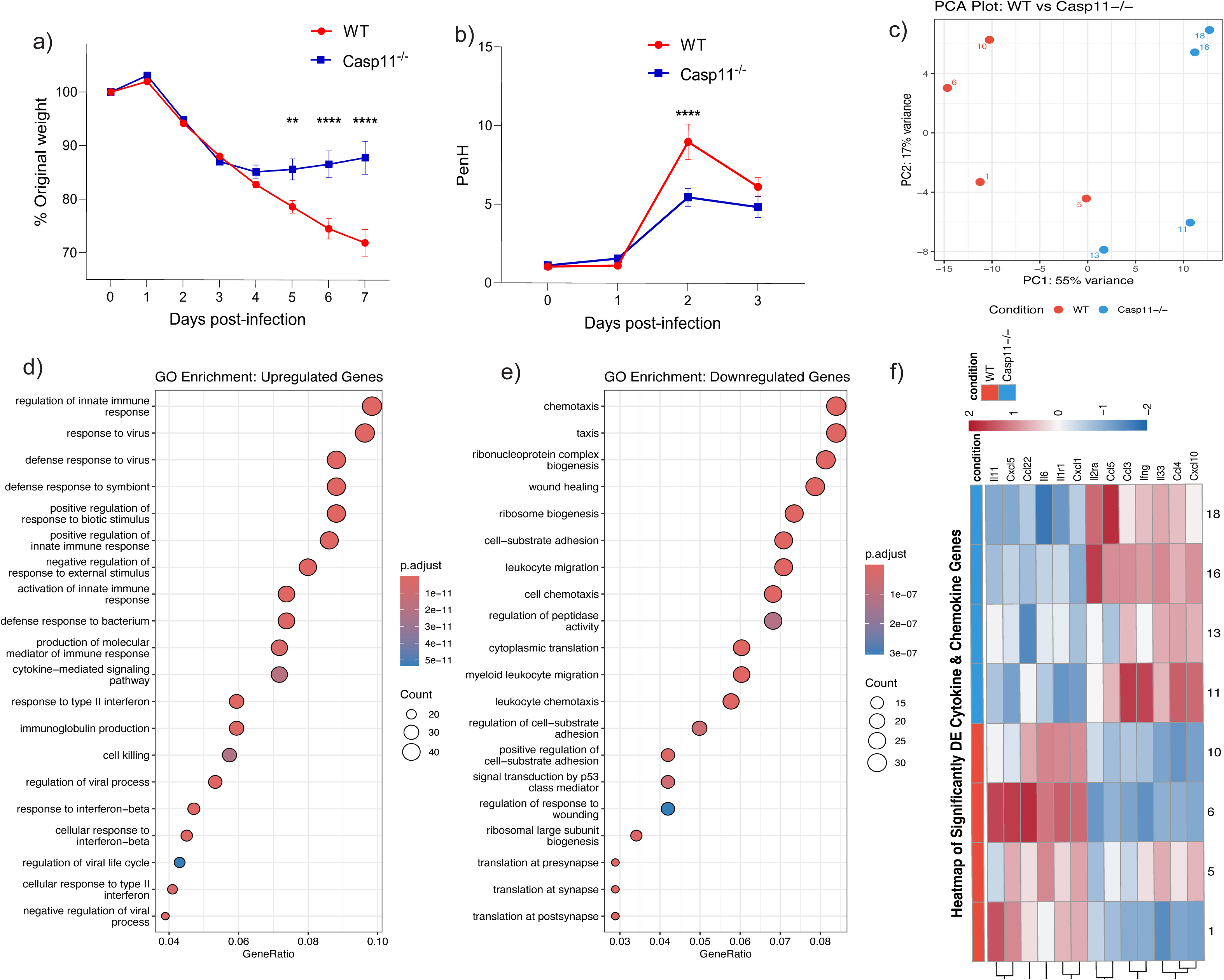
CASP11 deficiency protects against severe SARS⍰CoV⍰2 infection, limits early hyperinflammation and enhances interferon responses. (a) Body weight change in WT and *Casp11*⁻^/^⁻ mice following intranasal infection with 5*10^5 TCID50 of mouse-adapted (MA10) SARSl1CoVl12. WT mice exhibited progressive weight loss through day 7, whereas *Casp11*⁻^/^⁻ mice began recovering by day 4 (n = 14/group) (Equal number of males and females per group). Two-way ANOVA. (b) PenH, as a surrogate of airway resistance, measured by whole-body plethysmography at day 3 post-infection, showing significantly lower resistance in *Casp11*⁻^/^⁻ mice, indicative of improved lung function (n = 10/WT, 12/ *Casp11*⁻^/^⁻) (Equal number of males and females per group). Two-way ANOVA (c) Principal component analysis (PCA) of lung transcriptomes derived from bulk RNA-seq at day 4 post-infection shows clear separation between WT and *Casp11*⁻^/^⁻ samples, indicating genotype-driven differences in transcriptional responses (n = 4/group). Equal number of males and females per group. (d) Gene Ontology (GO) enrichment analysis of genes with higher gene expression in *Casp11*⁻^/^⁻ lungs compared to WT reveals enrichment in type I and II interferon signaling and antiviral immune pathways (n = 4/group). (e) GO enrichment analysis of genes with lower gene expression in *Casp11*⁻^/^⁻ lungs compared to WT highlights suppression of innate immune pathways, including myeloid cells chemotaxis and migration (n = 4/group). (f) Heatmap of differentially expressed cytokine and chemokine genes shows a shift in *Casp11*⁻^/^⁻ lungs from pro-myeloid inflammation (e.g., Cxcl1, Il6, Ccl22) to lymphocyte-promoting responses (e.g., Ccl3, Ccl5, Ifng) (n = 4/group). **p < 0.01; ****p < 0.0001.

To further investigate the molecular mechanisms underlying the enhanced protection and recovery observed in *Casp11*⁻^/^⁻ mice, we performed bulk RNA sequencing of lung tissue harvested at day 4 post-infection. Principal component analysis (PCA) revealed distinct transcriptional profiles between infected WT and *Casp11*⁻^/^⁻ lungs, highlighting a divergent host response **(Fig. 1c)**. Gene Ontology (GO) enrichment analysis of genes with higher expression in the *Casp11*⁻^/^⁻ group showed a marked enrichment of antiviral immune pathways. Notably, both type I and type II interferon response pathways were significantly upregulated, suggesting a robust antiviral state in the absence of CASP11**(Fig. 1d)**. In contrast, genes with lower expression in the *Casp11*⁻^/^⁻ lungs were predominantly related to myeloid cell chemotaxis and migration, suggesting a reduced innate inflammatory response in the absence of CASP11**(Fig. 1e)**.

Further gene set enrichment analysis (GSEA) revealed significant enrichment of antiviral pathways in *Casp11*⁻^/^⁻ lungs, including the interferon-γ (IFN-γ) response and lymphocyte-mediated immunity **(Suppl. Fig. 1)**. We then examined the expression of cytokines and chemokines in infected lungs. In *Casp11*⁻^/^⁻ mice, there was upregulation of IFNl1γ and several chemokines known to promote effector T cell recruitment, including CCL3, CCL4, CCL5, and CXCL10 (36,37) **(Fig. 1f) & (Suppl. Fig. 2).** In contrast, cytokines and chemokines typically associated with neutrophil and monocyte recruitment such as CXCL1, CXCL5, CCL22, IL-6, IL-11, and IL1R1 were downregulated in *Casp11*⁻^/^⁻ lungs relative to WT (38,39)(Fig. 1f), suggesting a shift away from pro-inflammatory innate immune signaling toward a more lymphocyte-driven antiviral response **(Fig. 1f) & (Suppl. Fig. 2)**.

We then examined cytokine protein levels in lung homogenates using ELISA at day 4 post-infection, the same time point used for the RNA-seq analysis. Compared to WT, *Casp11*⁻^/^⁻ lungs displayed markedly lower levels of ILl11β, ILl16, and CXCLl11, while interferonl1γ (IFNl1γ) levels were significantly elevated **(Fig. 2a)**. These findings confirm the transcriptional signature observed by RNA-seq and support a shift toward an upregulated interferon-γ response.

**Figure 2.**
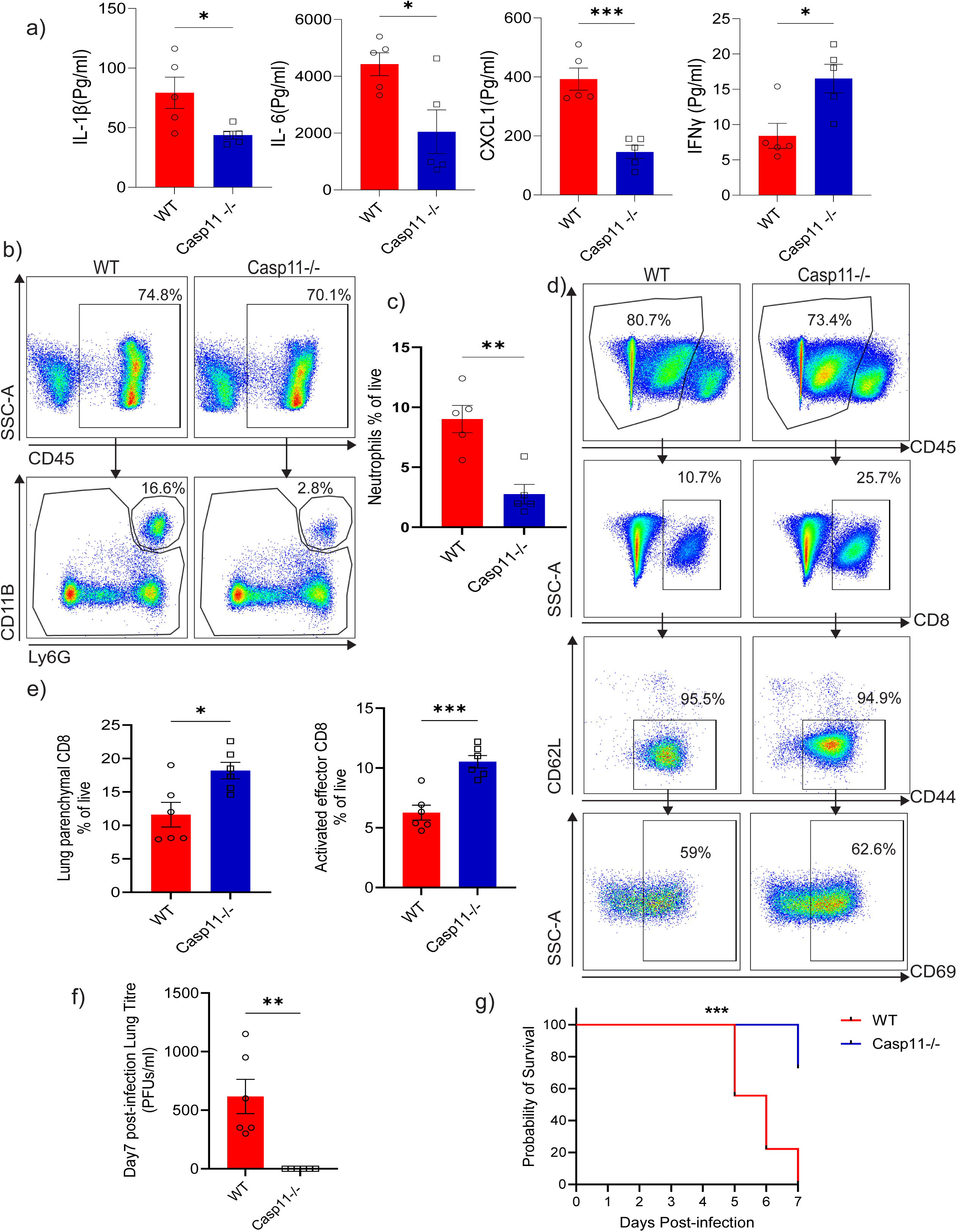
CASP11 deficiency limits early inflammation and neutrophil infiltration, enhances effector CD8⁺ T cell responses, promotes viral clearance, and protects against lethal SARS⍰CoV⍰2 infection. (a) Cytokine protein levels in lung homogenates at day 4 post-infection, measured by ELISA. *Casp11*⁻^/^⁻ lungs exhibited reduced levels of ILl11β, ILl16, and CXCLl11, and elevated IFNl1γ compared to WT, confirming RNA-seq findings (n = 5/group) (3 males & 2 females/group). Unpaired t-test. (b, c) Flow cytometry of lung tissue at day 4 post-infection shows significantly fewer neutrophils (CD45^hiCD11b^hiLy6G^hi) in *Casp11*⁻^/^⁻ mice compared to WT, consistent with reduced neutrophil-attracting chemokines. Gating strategy (b), Quantification (c), (n = 5/group) (3 males & 2 females/group). Unpaired t-test. (d, e) At day 7 post-infection, flow cytometry following intravascular anti-CD45 labeling revealed higher frequencies of lung parenchymal (CD45^lo) CD8⁺ T cells and activated effector CD8⁺ T cells (CD62L^loCD44^hiCD69^hi) in *Casp11*⁻^/^⁻ lungs. Gating strategy (d), Quantification (e), (n = 6/group) (Equal number of males and females per group). Unpaired t-test. (f) Viral titers in lung homogenates assessed by plaque assay at day 7 post-infection. *Casp11*⁻^/^⁻ mice completely cleared the virus by day 7, in contrast to WT (n = 6/group) (Equal number of males and females per group). unpaired t-test. (g) Kaplan–Meier survival analysis following high-dose SARSl1CoVl12 challenge (dose = 1*10^6 TCID50). All WT mice succumbed by day 7, while 72% of *Casp11*⁻^/^⁻ mice survived. (n = 6/group). Log-rank (Mantel-Cox) test. *p < 0.05; **p < 0.01; ***p < 0.001.

### CASP11 deficiency limits neutrophil infiltration, enhances effector CD8**⁺** T cell responses, promotes viral clearance, and protects against lethal SARS⍰CoV⍰2 infection

Given the distinct differences in immune responses between WT and *Casp11*⁻^/^⁻ mice, we next investigated the immune cell composition in the lungs following infection. Lungs were harvested on day 4 post-infection and analyzed by flow cytometry. Consistent with the observed downregulation of neutrophil-attracting chemokines, *Casp11*⁻^/^⁻ lungs exhibited significantly reduced neutrophil infiltration compared to WT-infected lungs **(Gating strategy Fig. 2b) (Fig. 2c)**, supporting a reduced innate inflammatory response in the absence of CASP11.

We then performed flow cytometry on lung tissue harvested at day 7 post-infection, a time point selected to allow for the development of an effector T cell response. To distinguish lung-resident (parenchymal) T cells from circulating peripheral T cells, mice were retro-orbitally injected with fluorescent anti-CD45 antibody 10 minutes before sacrifice, labeling CD45⁺ cells in the intravascular space while leaving tissue-cells (lung parenchymal cells) unlabeled. Peripheral blood cells were >90% CD45^hi, demonstrating efficient labeling of circulating cells (data not shown). Flow cytometric analysis revealed that *Casp11*⁻^/^⁻ lungs contained significantly more lung parenchymal CD8⁺ T cells compared to WT controls **(Gating strategy Fig. 2d) (Fig. 2e)**. Further analysis of T cell activation status, based on surface marker expression, demonstrated an increased proportion of activated effector CD8⁺ T cells (CD62L^loCD44^hiCD69^hi) in *Casp11*⁻^/^⁻ lungs relative to WT **(Gating strategy Fig. 2d) (Fig. 2e)**. No significant differences were observed in CD45^lo CD4⁺ T cell populations between the two groups **(Suppl. Fig. 3a)**.

To test whether this enhanced T cell response translates into improved viral clearance, we performed plaque assays on lungs collected from WT and *Casp11*⁻^/^⁻ mice at days 4 and 7 post-infection. While viral loads were similar between groups at day 4 (**Suppl. Fig. 3b)**, by day 7, *Casp11*⁻^/^⁻ mice had completely cleared the virus, in contrast to WT mice, which still had detectable viral titers **(Fig. 2f).** These results suggest that the robust activated effector CD8⁺ T cell response in *Casp11*⁻^/^⁻ mice is associated with an enhanced viral clearance.

We next asked whether CASP11 deficiency protects against lethal SARSl1CoVl12 infection. WT and *Casp11*⁻^/^⁻ mice were infected with a higher dose of MA10 (1*10^6 TCID50). All WT mice succumbed to infection by day 7, while 72% of the *Casp11*⁻^/^⁻ mice survived **(Fig. 2g)**. These data demonstrate that CASP11 deficiency promotes viral clearance and confers protection from lethal SARSl1CoVl12 challenge.

### CASP11 deficiency promotes functional, antigen-specific CD8**⁺** T cell response independent of SARS⍰CoV⍰2 disease severity

Next, we investigated whether the absence of CASP11 similarly promotes a robust effector CD8⁺ T cell response during moderate SARS-CoVl12 infection. To address this, we used a low-dose infection model in which WT and *Casp11*⁻^/^⁻ mice were infected with 1 × 10^5 TCID50 of the mouse-adapted SARS-CoVl12 strain MA10. As previously reported, both WT and *Casp11*⁻^/^⁻ mice began regaining weight by day 4 and continued to recover through day 7, although WT mice exhibited a slower recovery trajectory(32). *Casp11*⁻^/^⁻ mice also had reduced early inflammatory cytokines (ILl11β, ILl18, and ILl16) and decreased neutrophil infiltration at days 2 and 4 post-infection(32). Importantly, viral loads were comparable between strains, becoming nearly undetectable by day 4 and fully cleared by day 7(32).

To further evaluate the adaptive immune response in this less severe SARS-CoV-2 infection model, we performed flow cytometry on lung tissue at day 7 post-infection, a time point when both groups had cleared the virus. As in previous experiments, mice received a retro-orbital injection of fluorescent anti-CD45 antibody prior to euthanasia to label intravascular immune cells, enabling us to distinguish infiltrating lung parenchymal T cells.

We found that *Casp11*⁻^/^⁻ mice harbored a higher number of lung parenchymal CD8⁺ T cells compared to WT **(Gating strategy Fig. 3a) (Fig. 3b)**, including a significantly increased population of activated effector CD8⁺ T cells (CD62L^loCD44^hiCD69^hi) **(Gating strategy Fig. 3a) (Fig. 3c)**. These findings confirm that the enhanced CD8⁺ T cell response observed in *Casp11*⁻^/^⁻ mice is not merely a consequence of higher viral burden in WT mice. Instead, it is intrinsically driven by the absence of CASP11.

**Figure 3.**
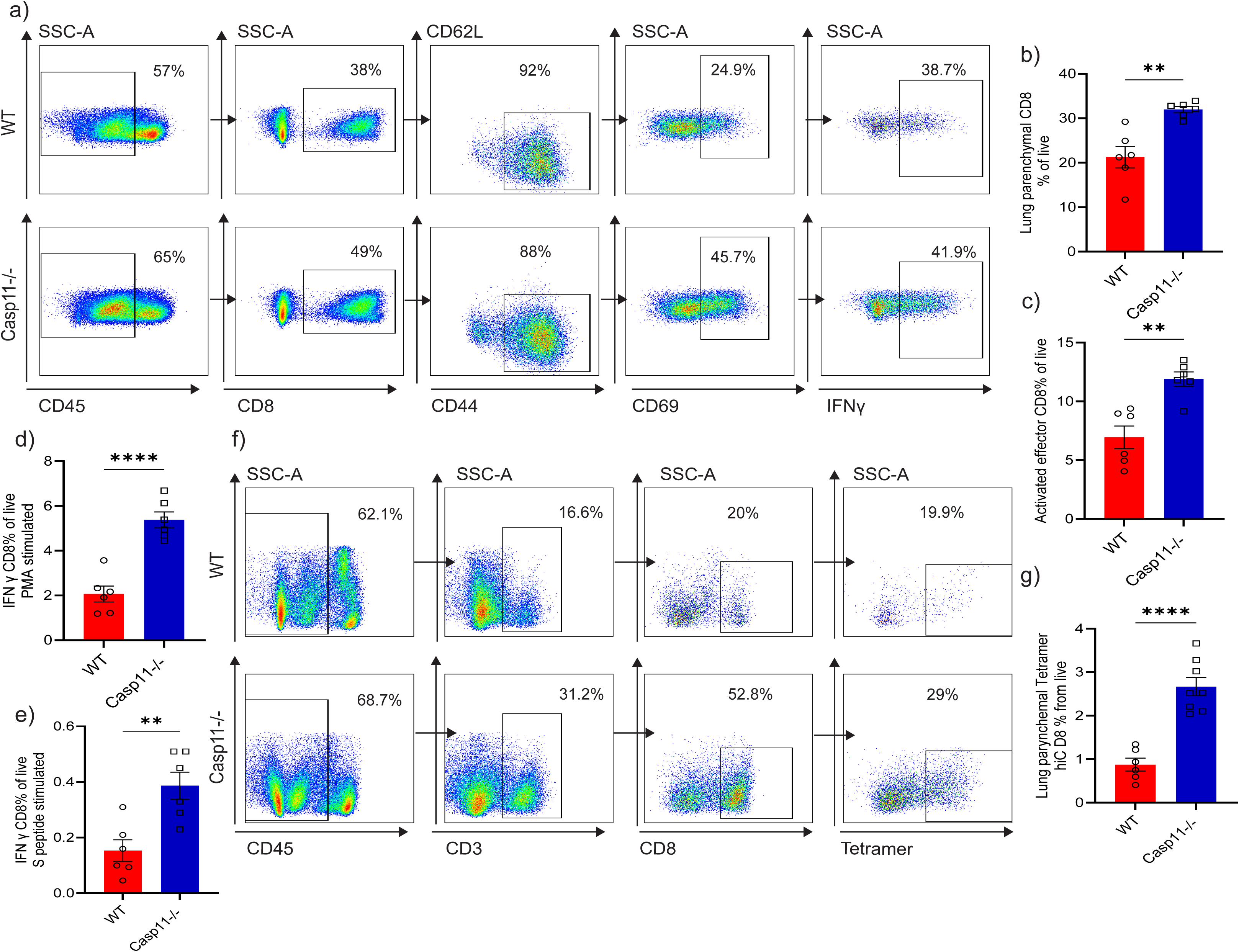
CASP11 deficiency promotes functional, antigen-specific CD8⁺ T cell responses independent of SARS⍰CoV⍰2 disease severity. (a-c) Representative flow cytometry plots showing lung parenchymal CD8⁺ T cells (CD45^loCD8^hi) and activated effector CD8⁺ T cells (CD62L^loCD44^hiCD69^hi) at day 7 post-infection in WT and *Casp11*⁻^/^⁻ mice following low-dose (1*10^5 TCID50) MA10 SARS-CoVl12 infection. Gating strategy (a), Quantification of lung parenchymal CD8⁺ T cells (b) and activated effector CD8⁺ T cells (CD62L^loCD44^hiCD69^hi) (c) as a percentage of live cells showing increased frequency of functional effector CD8⁺ T cells in *Casp11*⁻^/^⁻ lungs. (n= 6/group) (Equal number of males and females per group). Unpaired t-test. (d–e) Flow cytometry analysis of intracellular IFNl1γ production by CD8⁺ T cells after ex vivo stimulation, with PMA/Ionomycin (d) or SARSl1CoVl12 spike peptide (e), showing increased frequency of functional effector IFNl1γ CD8⁺ T cells in *Casp11*⁻^/^⁻ lungs. Gating strategy (a). (n= 6/group) (Equal number of males and females per group). Unpaired t-test. (f-g) Representative flow cytometry analysis of tetramer staining of lung CD8⁺ T cells using SARSl1CoVl12-specific tetramer (H-2D^b^ N_219-227_). Gating strategy (f), Quantification of tetramer positive CD8⁺ T cells among lung parenchymal T cells (g). *Casp11*⁻^/^⁻ mice exhibit a significantly greater population of antigen-specific CD8⁺ T cells. (n= 6/ WT, n=8/ *Casp11*⁻^/^⁻) (Equal number of males and females per group). Unpaired t-test. **p < 0.01; ****p < 0.0001.

Because CASP11 is a central effector in the inflammasome pathway, we also investigated whether its absence affects the T cell phenotype in response to SARS CoV-2 infection(40,41). To explore this, we isolated lung from WT and *Casp11*⁻^/^⁻ mice at day 7 post-infection and stimulated single cell suspensions ex-vivo with PMA/ionomycin or an overlapping peptide pool derived from SARSl1CoVl12 Spike protein. Following stimulation, cells were stained for T cell activation markers and intracellular IFNl1γ, since IFNl1γ producing CD8 T cells are generally critical for respiratory virus clearance and infection control. *Casp11*⁻^/^⁻ mice showed increased numbers of IFNl1γ–producing effector CD8⁺ T cells in response to both stimuli **(Gating strategy Fig. 3a) (Fig. 3d & Fig. 3e)**, suggesting that Casp11 deficiency promotes expansion or maintenance of an CD8⁺ T cell population with heightened effector function.

Furthermore, to evaluate the antigen specificity of the CD8⁺ T cell response, we performed SARSl1CoVl12 tetramer staining, which allows direct detection of CD8⁺ T cells specific for defined viral epitopes presented by MHC class I(42). This approach is especially useful for distinguishing true antigen-driven responses from bystander T cell activation, an important distinction in the context of viral infections like SARSl1CoVl12 infection. *Casp11*⁻^/^⁻ mice exhibited an increased number of lung parenchymal tetramer (H-2D^b^ N_219-227_) positive CD8⁺ T cells **(Gating strategy Fig. 3f) (Fig. 3g)**, indicating that the enhanced CD8 T cell repones detected in the absence of CASP11 is composed of virus antigen-specific effector cells. Overall, these findings demonstrate that CASP11 deficiency enhances the quantity and functional quality of virus-specific effector CD8⁺ T cells in the lung after SARS-CoV-2 infection.

### CASP11 mediates persistent inflammatory and T cell responses after SARS⍰CoV⍰2 infection, indicating ongoing inflammation and immune activation despite viral clearance

Persistent inflammation and sustained T cell activation have been implicated in post-viral sequelae, including tissue remodeling, immune dysregulation, and long-term complications such as fibrosis, impaired lung function, and symptoms contributing to long COVID(43–48). Therefore, we next investigated if CASP11 influences the persistence of inflammatory responses and immune cell activation after viral clearance, and whether it contributes to the prolonged immune activity often observed beyond the virus replication phase in SARS-CoV-2 infection. Given this, we aimed to determine whether CASP11 plays a role in maintaining inflammation and immune activation after the resolution of acute infection. To address this, we infected WT and *Casp11*⁻^/^⁻ mice with MA10(5x10^5 TCID50). Mice were then sacrificed at day 14, and lung tissue was collected for cytokine profiling and flow cytometric analysis. On day 14 post-infection, WT lungs exhibited persistently elevated levels of inflammatory cytokines (**Suppl Fig. 4a)** and a higher number of CD8⁺ and CD4⁺ T cells than the lungs of *Casp11*⁻^/^⁻ mice **(Suppl Fig. 4b & 4c)**. In contrast, *Casp11*⁻^/^⁻ lungs demonstrated a more resolved immune state, characterized by lower cytokine levels and reduced T cell number. These findings suggest that CASP11 promotes prolonged post-viral inflammation and immune activation, potentially contributing to delayed immune resolution and long-term tissue damage following SARSl1CoVl12 infection.

### CASP11 deficiency in hematopoietic immune cells enhances survival, reduces inflammation, and promote T cell response after SARS⍰CoV⍰2 infection

To further elucidate the mechanism by which CASP11 deficiency enhances the adaptive immune response and protects against severe disease, we generated reciprocal bone marrow (BM) chimeric mice **(Suppl Fig. 5a)**. In this model, lethally irradiated WT recipient mice were reconstituted with bone marrow from either WT or *Casp11*⁻^/^⁻ donors, generating two groups: WT → WT chimeras (WT chimera) and *Casp11*⁻^/^⁻→ WT chimeras (hereafter referred to as *Casp11*⁻^/^⁻ chimeras). This strategy enabled us to isolate the role of CASP11 within radiosensitive hematopoietic cells, such as monocytes, dendritic cells, and lymphocytes, while preserving CASP11 expression in radioresistant non-hematopoietic compartments, including lung epithelial and endothelial cells. Using this model, we assessed the contribution of hematopoietic CASP11 to SARS-CoV-2 induced immunopathology, T cell responses, and viral clearance. In this model, interstitial macrophages and monocyte-derived macrophages in the lung are donor-derived, whereas alveolar macrophages are largely host-derived and remain WT due to their radioresistant, self-renewing nature(49). This setup enables a compartment-specific interpretation of CASP11 function in the lung immune microenvironment.

After a full reconstitution period, chimeric mice were infected with 1 × 10^5 TCID50 of MA10. This dose, in the context of relatively aged chimeras (approximately 6-8 months old at infection), consistently induced severe disease. Only 41% in the WT chimeras survived by 7 days post-infection, compared to 76% in the *Casp11*⁻^/^⁻ chimeras **(Fig. 4a)**. Respiratory functions in infected mice were assessed by plethysmography, which revealed that *Casp11*⁻^/^⁻ infected chimera mice had a decreased PenH values indicative of decreased airway resistance compared to infected WT chimeras **(Fig. 4b).** This indicates that the deficiency of CASP11 in hematopoietic cells significantly alters lung functions and clinical severity in SARS-Cov-2 infection. Consistent with these findings, H&E staining of lung sections demonstrated that WT chimeras had significantly more lung consolidation, with increased edema, inflammation, and cellular infiltrates at day 4 postl1infection compared to *Casp11*⁻^/^⁻ chimeras (**Fig. 4c & 4d**). Flow cytometric profiling of lung myeloid cells at day 4 post-infection showed fewer infiltrating neutrophils in *Casp11*⁻^/^⁻ chimeras compared to WT **(Fig. 4e**).

**Figure 4.**
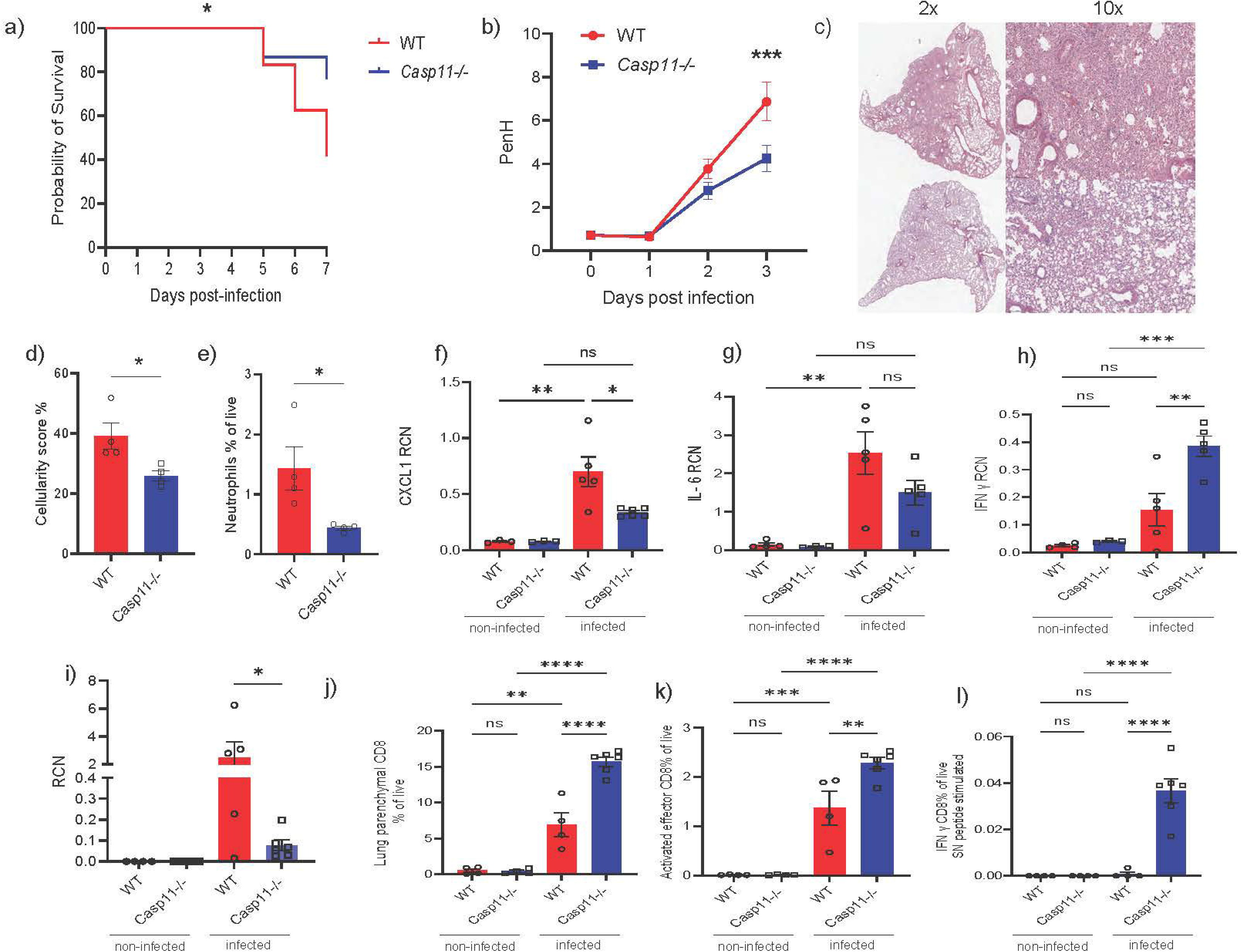
CASP11 deficiency in hematopoietic immune cells enhances survival, reduces inflammation, and promote T cell response after SARS⍰CoV⍰2 infection. (a) Kaplan–Meier survival analysis of WT → WT (WT chimeras) and *Casp11*⁻^/^⁻ → WT (*Casp11*⁻^/^⁻ chimeras) mice following infection with 1*10^5 TCID50 of MA10 SARSl1CoVl12. Casp11⁻/⁻ chimeras exhibited significantly improved survival. Only 41% in the WT chimeras survived by 7 days post-infection, compared to 76% in the *Casp11*⁻^/^⁻chimeras. (n = 24/ WT, n= 30/ *Casp11*⁻^/^⁻). Log-rank (Mantel-Cox) test. (b) PenH, as a surrogate of airway resistance, measured by whole-body plethysmography at day 3 post-infection, showing significantly lower resistance in *Casp11*⁻^/^⁻ chimera mice, indicative of improved lung function. (n= 18/ group). Two-way ANOVA (c–d) Representative H&E-stained lung sections at 2x and 10x magnification(c) and corresponding cellularity scores (d) at day 4 post-infection demonstrate reduced lung inflammation and edema in *Casp11*⁻^/^⁻ chimeras. (n= 4/ group). unpaired t-test. (e) Flow cytometric quantification of neutrophils in lung tissue at day 4 post-infection, showing reduced neutrophil infiltration in *Casp11*⁻^/^⁻ chimeras. (n= 4/ group). unpaired t-test. (f–h) RT-qPCR analysis of inflammatory cytokines and chemokines in lung tissue shows downregulation of Cxcl1 (f) and Il6(g), and upregulation of Ifng (h) in *Casp11*⁻^/^⁻chimeras. (n = 5/ WT, n= 6/ *Casp11*⁻^/^⁻ ). Two-way ANOVA (i) Viral RNA copy number in lung homogenates at day 7 post-infection, showing significantly reduced viral loads in *Casp11*⁻^/^⁻ chimeras. (n = 5/ WT, n= 6/ *Casp11*⁻^/^⁻). unpaired t-test. (j–k) Flow cytometric quantification of lung parenchymal CD8⁺ T cells (j) and activated effector CD8⁺ T cells (CD62L^loCD44^hiCD69^hi) (k) at day 7 post-infection. (n = 4/ WT, n= 6/ *Casp11*⁻^/^⁻). Two-way ANOVA (l) Intracellular staining of IFNl1γ⁺ CD8⁺ T cells following ex vivo stimulation with spike+nucleocapsid (SN) peptide mixture, indicating enhanced effector function in *Casp11*⁻^/^⁻ chimeras. (n = 4/ WT, n= 6/ *Casp11*⁻^/^⁻ ). Two-way ANOVA *p < 0.05; **p < 0.01; ***p < 0.001; ****p < 0.0001; ns, not significant.

Cytokine mRNA analysis by qPCR revealed that ILl16 and CXCL1 were downregulated, while IFNl1γ was significantly upregulated in Casp11⁻/⁻ chimera mice **(Fig. 4f-4h)**, consistent with earlier observations in global knockouts. To investigate broader transcriptional changes, we performed RNA-seq on lung tissue at day 4. Gene set enrichment analysis (GSEA) showed that Casp11⁻/⁻ chimeras had higher gene expressions related to T cell activation and antigen presentation pathways **(Suppl Fig. 5b)**.

By day 7 post-infection, *Casp11*⁻^/^⁻ chimeras exhibited significantly lower viral copy numbers in the lungs compared to WT chimeras, indicating enhanced viral clearance **(Fig. 4i)**. To assess T cell response, we performed in vivo labeling of intravascular immune cells via retro-orbital injection of fluorescent anti-CD45 antibody, as previously described. This allowed us to selectively quantify lung parenchymal CD8⁺ T cells. Flow cytometric analysis revealed a higher number of parenchymal CD8⁺ T cells in *Casp11*⁻^/^⁻ chimeras relative to WT controls **(Fig. 4j)**, along with a greater frequency of activated effector CD8⁺ T cells characterized by a CD62L^loCD44^hiCD69^hi phenotype **(Fig. 4k)**.

To further evaluate CD8⁺ T cell phenotype effector function, we stimulated lung-derived CD8⁺ T cells ex vivo with a SARS-CoV-2 Spike and Nucleocapsid peptide (SN peptide) mixture. *Casp11*⁻^/^⁻ chimeras demonstrated a higher frequency of IFNl1γ producing CD8⁺ T cells in response to both stimuli, indicating preservation of effector function and enhanced antiviral potential **(Fig. 4l) (Gating strategy Suppl Fig. 6)**.

### Mononuclear phagocyte system (MPS) cell-specific CASP11 deletion enhances antiviral CD8**⁺** T cell responses, accelerates recovery, and improves viral clearance during SARS⍰CoV⍰2 infection

The enhanced adaptive immune response observed in *Casp11*⁻^/^⁻ chimeras, along with reduced disease severity and improved viral clearance, suggests that CASP11 exerts its effects through hematopoietic immune cells. However, it remains unclear whether this regulation is primarily mediated by myeloid cells via modulation of the innate immune response, or whether CASP11 also plays a direct, intrinsic role in T cells.

To address this question, we generated mice with CASP11 deficiency specifically in mononuclear phagocyte system (MPS) cells. We utilized Cx3cr1l1CreERT mice to generate Casp11-deficient mice in CX3CR1-expressing lineages, predominantly targeting the mononuclear phagocyte system (MPS)(50). The main cell types affected by this system include monocytes, monocyte-derived macrophages, interstitial macrophages, and some lineages of dendritic cells(50). Alveolar macrophages are minimally impacted due to their low expression of CX3CR1(50,51). Also, neutrophils and lymphocytes do not express CX3CR1, or they do so at very low to undetectable levels, both in mice and humans (50,52). Therefore, they are not significantly affected by CX3CR1-CreERT2–mediated recombination(50,52). For simplicity, we refer to Cx3cr1l1CreERT mice as "WT" and Cx3cr1l1CreERT CASP11 flox mice as " *Casp11*⁻^/^⁻flox"

Both groups were infected with 5*10^5 TCID50 of MA10 virus to establish a severe SARSl1CoVl12 infection model. *Casp11*⁻^/^⁻ mice exhibited enhanced recovery, indicating milder disease severity (Fig. 5a). Histological analysis (H&E staining) and cellularity scoring of lung sections at day 4 post-infection revealed reduced inflammatory infiltrates and edema in *Casp11*⁻^/^⁻ mice compared to WT **(Fig. 5b & 5c)**.

**Figure 5.**
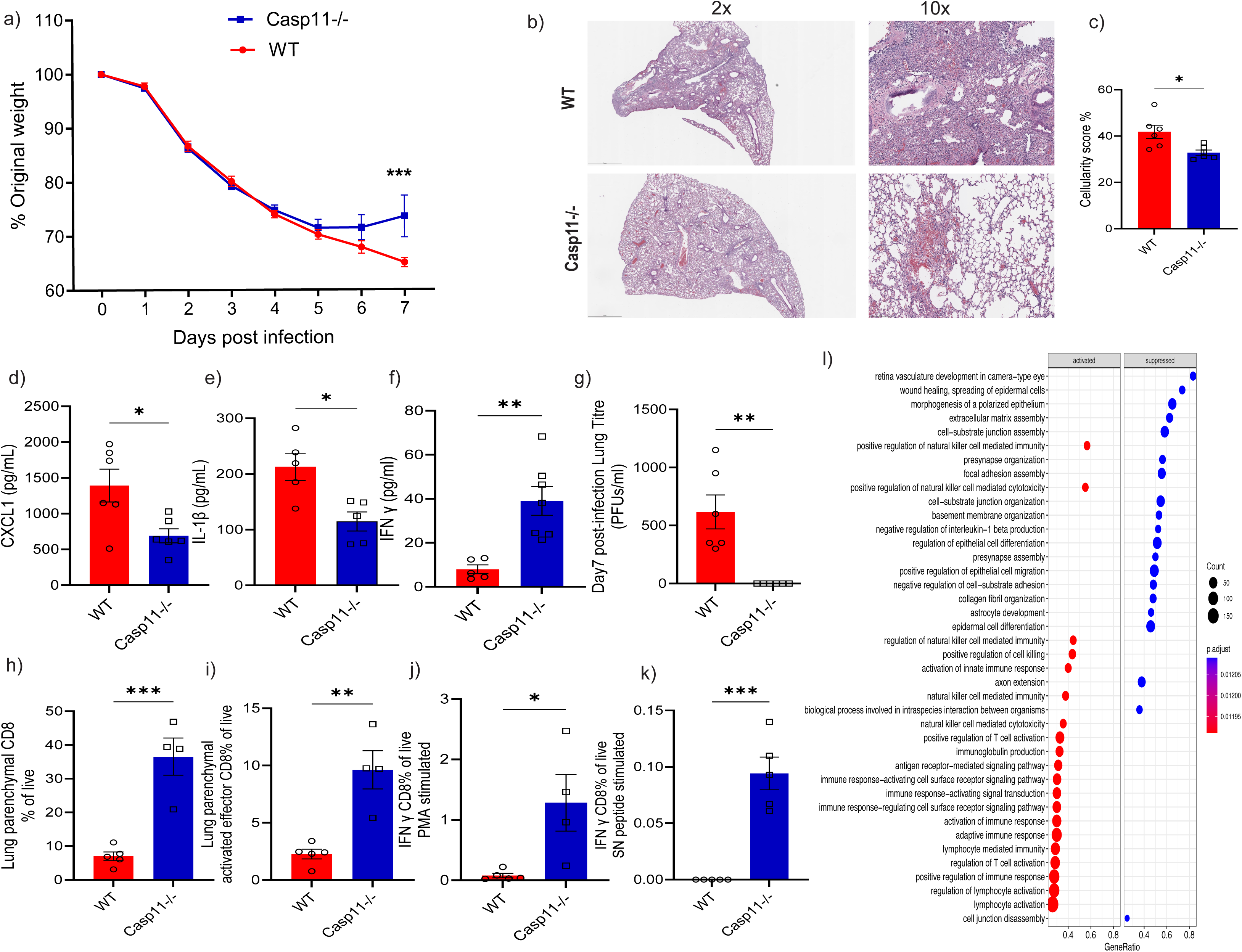
Mononuclear phagocyte system (MPS) cell-specific Caspase⍰11 deletion enhances antiviral CD8⁺ T cell responses, accelerates recovery, and improves viral clearance during SARS⍰CoV⍰2 infection. (a) Weight loss kinetics in WT and *Casp11*⁻^/^⁻ flox mice following intranasal infection with 5*10^5 TCID50 of MA10 SARSl1CoVl12. Casp11⁻/⁻ flox mice showed significantly improved recovery. (n = 11/ group). Log-rank (Mantel-Cox) test. (b) Representative H&E-stained lung sections at day 4 post-infection show reduced inflammatory infiltrates in *Casp11*⁻^/^⁻ flox mice compared to WT mice. 2x and 10x magnification. (c) Histopathologic cellularity scoring of lung sections confirms reduced inflammation in *Casp11*⁻^/^⁻ flox mice. (n = 5/ group). unpaired t-test. (d–f) Cytokine analysis from lung homogenates at day 4 post-infection shows reduced IL-1β (d) and CXCL1 (e), and elevated IFN-γ (f) in *Casp11*⁻^/^⁻ flox mice compared to WT. (n = 6/ WT, n= 7/ *Casp11*⁻^/^⁻). unpaired t-test. (g) Plaque assay quantification of viral titers in lung tissue at day 7 post-infection reveals significantly lower viral loads in *Casp11*⁻^/^⁻ flox mice. (n=6/ group). unpaired t-test. (h) Quantification of lung parenchymal CD8⁺ T cells (CD45⁻ CD8⁺) by flow cytometry at day 7 post-infection shows increased numbers in *Casp11*⁻^/^⁻ flox mice. (n=5/ group). unpaired t-test. (i) Frequency of activated effector CD8⁺ T cells (CD62L^loCD44^hiCD69^hi) among lung parenchymal CD8⁺ T cells. (n=5/ group). unpaired t-test. (j–k) Frequency of IFNl1γ⁺ CD8⁺ T cells following ex vivo stimulation with PMA (j) or spike+nucleocapsid (SN) peptide mix (k) shows increased cytokine production in *Casp11*⁻^/^⁻flox mice. (n=5/ group). unpaired t-test. (l) Gene Set Enrichment Analysis (GSEA) of lung transcriptomes at day 4 reveals enrichment of antigen presentation, T cell activation, and adaptive immune response pathways in *Casp11*⁻^/^⁻ flox mice relative to Cre controls. (n=5/ group). *p < 0.05; **p < 0.01; ***p < 0.001.

We next investigated the cytokine release profile in lung tissue harvested at day 4 post-infection. *Casp11*⁻^/^⁻ mice showed reduced release of inflammatory cytokines, including IL-1β and CXCL1 **(Fig. 5d & 5e)**, while exhibiting elevated levels of IFN-γ **(Fig. 5f)**, consistent with observations in global *Casp11*⁻^/^⁻ knockout and chimera mice. These findings confirm that modulation of the early innate immune response and the resulting lung protection is mediated mainly by CASP11 within MPS cells.

We then used plaque assay to quantify the viral load in the lungs of infected mice at day 7 post-infection. We found that *Casp11*⁻^/^⁻ mice had significantly lower viral loads compared to WT mice, indicating that the enhanced viral clearance observed in global *Casp11*⁻^/^⁻ knockouts and chimeras is due to the immunomodulatory effect of CASP11 in monocyte derived cells **(Fig. 5g)**.

Next, we investigated changes in the CD8⁺ T cell response mediated by Casp11 deficiency in monocyte cells. Cre and *Casp11*⁻^/^⁻ flox mice were infected with the same dose of MA10 (2*10^5 TCID50) and sacrificed at day 7 post-infection, after retro-orbital injection of fluorescent anti-CD45 antibody, as described before. Single-cell suspensions were prepared from the lungs, stained for surface markers, and subsequently stimulated ex vivo with PMA and a mixture of spike and nucleocapsid (SN) peptides, followed by permeabilization and intracellular interferon-γ staining. Flow cytometric analysis revealed that *Casp11*⁻^/^⁻ flox mice had a higher number of lung parenchymal CD8⁺ T cells compared to WT mice **(Fig. 5h)**. Additionally, analysis of CD69⁺ activated effector CD8⁺ T cells showed significantly increased numbers in *Casp11*⁻^/^⁻ flox mice **(fig. 5i)**.

Intracellular cytokine staining demonstrated a higher frequency of IFN-γ–producing CD8⁺ T cells in the lungs of *Casp11*⁻^/^⁻ flox mice compared to WT mice following stimulation with both PMA and SN peptide mixtures **(Fig. 5j & 5k)**. Moreover, bulk RNA sequencing of lung tissue at day 4 post-infection, coupled with gene set enrichment analysis (GSEA), revealed enhanced antigen presentation and upregulation of lymphocyte activation and T cell activation pathways **(Fig. 5l)**. Collectively, these findings demonstrate that the upregulated interferon-γ response and the more robust adaptive CD8⁺ T cell response, leading to accelerated recovery and viral clearance, are mediated by the absence of CASP11 in myeloid cells, primarily lung interstitial macrophages, circulating monocytes and monocyte-derived macrophages.

## Discussion

The absence of effective therapies addressing both viral replication and immune dysregulation remains a significant clinical gap in SARS-CoV-2 acute and post-acute conditions. Although previous clinical reports have documented suboptimal and delayed adaptive Tl1cell responses in severe SARS-CoV-2 infection patients, the underlying mechanisms driving these impaired Tl1cell responses remains unclear (12–19). While much attention has been given to the hyperinflammatory responses driving severe SARS-CoV-2 infection, the mechanisms by which this dysregulation undermines antiviral T cell immunity remain poorly understood (5,14,53–59).

Our findings suggest that targeting CASP11 can effectively address this gap. CASP11 deficiency enhances early tissue-resident effector CD8⁺ T cell responses, without functional impairments in T cells, such as altered phenotype, diminished cytokine production, or compromised antigen specificity leading to protection from severe SARS-CoV-2 infection and more efficient viral clearance. These results uncover an unexpected role for CASP11 in impairing adaptive immunity, extending its known function beyond innate immune activation and pyroptosis. CASP11 deficiency shifts lung immunity from innate hyperinflammation toward an enhanced, early, protective adaptive T-cell response, leading to more efficient viral clearance. Specifically, *Casp11*⁻^/^⁻ mice exhibited a robust interferon-gamma (IFN-γ) response with enhanced expression of T-cell recruiting chemokines (CCL3, CCL4, CCL5, and CXCL10), which facilitated CD8 T-cell infiltration and/or proliferation in infected lungs.

Mounting a robust interferon (IFN) response early during SARS-CoV-2 infection is crucial for establishing effective antiviral immunity, as impaired IFN signaling has been detected in patients with severe SARS-CoV-2 infection (60–62). Moreover, several studies have reported diminished and delayed IFN-γ responses in severe SARS-CoV-2 infection, correlated strongly with impaired T-cell functionality, increased disease severity, and viral persistence(63–66). Our findings align with and expand upon these reports, demonstrating that targeting CASP11 is a promising strategy to enhance IFN-γ response, which rapidly facilitates a shift toward an effective antiviral CD8 T-cell-mediated response. IFN-γ is critically involved in the upregulation of chemokines responsible for recruiting and activating T cells like CXCL10, as well as directly skewing immune responses towards CD8 cytotoxic T cells(67–69). *Casp11*⁻^/^⁻ mice also exhibited a potent T-cell favoring chemokine signature, including upregulated expression of CCL3, CCL4, CCL5, and CXCL10(36,37). These chemokines are known to interact with the chemokine receptors, primarily expressed on CD8 T cells, thereby promoting efficient T-cell recruitment, proliferation, and differentiation into tissue-resident memory (TRM) cells. This provides a mechanistic link between innate immune response, enhanced IFN-γ signaling, chemokine-driven CD8 T-cell responses, and improved viral clearance mediated by the absence of CASP11.

*Casp11*⁻^/^⁻ mice also exhibited a more robust type I interferon response early during infection. While this enhanced interferon signaling may contribute to the initial containment of viral replication, it is unlikely to fully account for the improved viral clearance observed later in infection. Notably, both WT and *Casp11*⁻^/^⁻mice exhibited comparable viral loads at day 4 post-infection, yet by day 7, *Casp11*⁻^/^⁻ mice showed a marked reduction in lung viral titers. This timing strongly correlates with the expansion and activation of virus-specific CD8⁺ T cells, suggesting that adaptive immunity, rather than innate viral control alone, is the primary driver of improved viral clearance in the absence of CASP11. Multiple studies have demonstrated that T cells rather than Type I IFN are essential for SARS-CoVl12 clearance in mouse models(70–72). *Rag1*⁻^/^⁻ mice, which lack functional T and B cells, fail to control SARS-CoVl12 replication and exhibit persistent viral loads despite intact innate responses(70,71). These findings align with our data, supporting that CASP11 deficiency facilitates viral clearance primarily by enhancing CD8⁺ T cell–mediated responses.

Although the exact mechanism by which the absence of CASP11 modulates the adaptive immune is unclear, it appears to be multifaceted. It can be an interplay between upregulating the interferon response and suppressing the neutrophil and proinflammatory cytokine response. Suppression of neutrophil infiltration, an indicator of severe inflammation, is known to impair antiviral T-cell responses. In our model, *Casp11*⁻^/^⁻ mice showed significantly reduced neutrophil infiltration through day 4 post-infection compared to WT controls. We previously demonstrated that CASP11 is also required for neutrophil extracellular trap (NET) formation during SARS-CoV-2 infection(32). This is particularly relevant because persistently elevated neutrophils and excessive NETs have been shown to hinder T-cell activation and proliferation(73). Additionally, CASP11 dependent inflammatory cytokines such as ILl11β, ILl16, and ILl18 may contribute to further delays in T-cell responses(74). Notably, *Casp11*⁻^/^⁻ mice exhibited lower levels of these proinflammatory cytokines in the lungs during severe SARS-CoV-2 infection.

CASP11 deficiency and the consequent early recruitment of activated effector CD8 T cells effectively controlled severe infection and significantly improved viral clearance. Even under conditions of high viral load intended to induce lethal disease, CASP11 deficiency markedly promoted survival, highlighting the therapeutic potential of specifically targeting CASP11. This contrasts with the published results of targeting other inflammasome components, such as Caspasel11 or NLRP3, which resulted in anti-inflammatory effect and a reduction of disease severity but failed to control viral replication (27). Current available therapeutic interventions for severe SARS-CoV-2 infection primarily depend on immunosuppressive agents, such as corticosteroids and ILl16 inhibitors, which mitigate hyperinflammatory responses without directly promoting antiviral immunity or enhancing viral clearance(75–77). Therefore, CASP11 targeting represents a disease-modifying therapeutic strategy rather than merely an immunosuppressive approach.

While a robust T-cell response is essential for effective viral clearance, accumulating evidence suggests that persistent T-cell activation may contribute to SARS-CoV-2 post-viral pathology. In mouse models, the T cell response to SARS-CoVl12 was associated with increased lung inflammation and injury, despite improved viral clearance(70). Recent clinical studies have reported that individuals with long-term respiratory complications or long COVID exhibit prolonged SARS-CoV-2-specific T-cell activation even months after viral clearance, in contrast to those who recover fully. This sustained immune activation has been implicated in chronic inflammation, impaired lung function, and tissue remodeling (43–48). Infected *Casp11*⁻^/^⁻ mice displayed resolution of inflammatory response at day 14 post infection with no detectable cytokine production and less inflammatory T cells. In contrast, WT mice exhibited persistent pro-inflammatory cytokine expression at day 14 and sustained activation of CD4⁺ and CD8⁺ T cells despite complete viral clearance. These findings suggest that CASP11 not only amplifies early inflammation but also impairs immune resolution, thereby contributing to prolonged immune activation and potential long-term tissue damage.

To further dissect the compartment-specific role of CASP11, we employed a reciprocal bone marrow chimera model. By transplanting bone marrow from *Casp11*⁻^/^⁻ or WT donors into lethally irradiated WT hosts, we created mice in which only hematopoietic-derived cells lacked CASP11, while radioresistant structural cells, including epithelial, endothelial, and alveolar macrophages, retained WT expression. In this setting, *Casp11*⁻^/^⁻ chimeras recapitulated the protective phenotype seen in global knockouts, with enhanced survival, improved lung function, reduced inflammation, and more robust adaptive immunity following SARSl1CoVl12 infection. Using this system, we demonstrated that CASP11 functions primarily within peripheral hematopoietic cells, underscoring their central role in shaping the immune response and determining disease outcomes after SARS-CoV-2 infection. Yet, it remained unclear how CASP11, a molecule traditionally associated with innate immunity, could influence adaptive T-cell responses and viral clearance. While prior studies have hinted at intrinsic roles for CASP11 in CD8⁺ T cell biology, we resolved this question by employing Cx3cr1l1CreERT Casp11^flox/flox mice, which selectively delete Casp11 in CX3CR1-expressing MPS cells, including monocytes, interstitial macrophages, and dendritic cells. This allowed us to isolate the effect of CASP11 in monocyte derived cells without affecting the expression of CASP11 in lymphoid or structural compartments. Strikingly, even with CASP11 deletion confined to monocyte cells, we observed an upregulated IFN-γ response, pronounced enhancement of CD8⁺ T cell activation, increased lung parenchymal CD8⁺ T cells, and a greater frequency of IFN-γ–producing effector cells. These findings argue against a cell-intrinsic role for CASP11 within CD8⁺ T cells and instead suggest that its immunomodulatory effects in SARS-CoV-2 are mediated through shaping the innate immune microenvironment.

We demonstrate, for the first time, that targeting CASP11 in monocyte modulates the innate immune environment in a way that protects against severe SARS-CoV-2 infection while also enhancing the development of a robust CD8⁺ adaptive immune response, ultimately leading to more efficient viral clearance. These findings suggest that CASP11 acts as an immunomodulatory molecule, capable of dampening harmful inflammation without broadly suppressing host immunity, distinguishing it from current immunosuppressive interventions used in severe SARS-CoV-2 infection.

## Methods

### Biosafety

Most experiments involving live SARS-CoV-2 were conducted in the OSU BSL-3 biocontainment facility. All procedures were reviewed and approved by the OSU BSL-3 Operations/Advisory Group, the Institutional Biosafety Officer, and the Institutional Biosafety Committee. Following the reclassification of SARS-CoV-2 to BSL-2, subsequent experiments were performed in a BSL-2 biocontainment facility in accordance with OSU biosafety guidelines.

### Viruses and titers

Mouse adapted SARS-CoV-2, variant strain MA10 (35), generated by the laboratory of Dr. Ralph Baric (University of North Carolina) was provided by BEI Resources (Cat # NR-55329). Viral stocks from BEI Resources were plaque purified on Vero E6 cells to identify plaques lacking mutations in the polybasic cleavage site of the Spike protein via sequencing. Non-mutated clones were propagated on Vero E6 cells stably expressing TMPRSS2 (provided by Dr. Shan-Lu Liu, The Ohio State University). Virus aliquots were flash frozen in liquid nitrogen and stored at -80 C. Virus stocks were sequenced to confirm a lack of tissue culture adaptation in the polybasic cleavage site. Virus stocks and tissue homogenates were titered on Vero E6 TMPRSS2 cells.

### Mice

C57BL/6 wild-type (WT) mice were obtained from The Jackson Laboratory (Bar Harbor, ME, USA). *Casp11*⁻^/^⁻ mice were generously provided by Dr. Yuan (Harvard Medical School, Boston, MA, USA). CX3CR1-CreER mice (JAX stock no. 021160) were obtained from The Jackson Laboratory. Casp11^flox/flox mice were kindly provided by Dr. Timothy Millar (University of Pittsburgh, Pittsburgh, PA, USA). Experiments were conducted with approval from the Animal Care and Use Committee at the Ohio State University (Columbus, OH, USA) which is accredited by AAALAC International according to guidelines of the Public Health Service as issued in the Guide for the Care and Use of Laboratory Animals.

### Sex as a biological variable

Our infection studies utilized both male and female animals, and we observed similar outcomes for both male and female mice. Pooled male and female data are shown throughout the manuscript.

### Chimera Mice

Reciprocal bone marrow (BM) chimeric mice with Casp11-/- or WT BM cell donors and WT irradiated hosts. *Casp11*⁻^/^⁻ → WT chimeras and WT → WT chimeras were constructed. BM chimeras will be constructed as described before(78). Recipient mice were lethally irradiated with two doses of 6.5 Gy and reconstituted by tail vein injection of 2 × 10^6 to 4 × 10^6 donor BM cells. Recipient were allowed to reconstitute their circulating leukocyte pools for six weeks prior to infection.

### CD45 cell labeling

To distinguish between lung parenchymal and intravascular T cell populations, anti-CD45-PE (clone 30-F11, BD Biosciences; 3 μg in 100 μL sterile PBS) was retro-orbitally injected 10 minutes prior to euthanasia to label circulating lymphocytes, while resident lung parenchymal lymphocytes remained unlabeled (7, 8). Peripheral blood collected at the time of sacrifice confirmed >90% labeling of circulating lymphocytes by flow cytometry.

### Tissue dissociation and flow cytometry

Lungs were dissected into single lobes before being dissociated into single cell suspension using gentleMACS octo-dissociator and Miltenyi lung dissociation kit (Miltenyi Biotec, 130-095-927). Red blood cells (RBCs) were lysed by incubating cells in 2 ml ACK buffer for 5 min at room temperature. After RBCs lysis, cells were washed in DPBS containing 1% BSA. The single cell suspension was centrifuged, and the cell pellets were washed twice with PBS. Cells were resuspended in T cell medium (RPMI 1640 supplemented with 0.1% gentamicin, 10% heat-inactivated FBS, Glutamax, and 5 × 10⁻ M β-ME) and incubated for 4–5 h at 37 °C in the presence of a protein transport inhibitor cocktail (eBioscience). Stimulation was performed either nonspecifically with PMA (50 ng/mL) and ionomycin (500 ng/mL), or with SARS-CoV-2 peptide pools (Miltenyi Biotec) at 1 µg/mL per peptide. These included PepTivator® Prot_S complete, Prot_S B.1.1.529/Omicron BA.1, Prot_S B.1.1.529/Omicron BA.2 (Refs. 130-127-951, 130-129-928, 130-130-807), and when indicated, PepTivator® Prot_N (Ref. 130-126-698). Cells incubated with DMSO alone served as negative controls. For some experiments, a combination of Spike and Nucleocapsid peptides was used for stimulation.

Following stimulation, cells were washed with cold PBS and stained with Live/Dead Zombie NIR Fixable Viability Dye (BioLegend, Cat. 423105) or LIVE/DEAD™ Fixable Aqua Dead Cell Stain Kit (Invitrogen, Cat. L34957) for 30 min at 4 °C. After two washes in PBS with 1% FBS (FACS buffer), cells were resuspended in Fc Block (clone 93; eBioscience, Cat. 14-0161-86) for 5 min at 4 °C, then stained with a combination containing the following surface antibodies for 20 min at 4 °C: CD3 V450 (clone 17A2; BD Biosciences, Cat. 561389), CD4 BV750 (clone H129.19; BD Biosciences, Cat. 747275), CD44 PerCP-Cy5.5 (clone IM7; BD Biosciences, Cat. 560570), CD62L BV605 (clone MEL-14; BD Biosciences, Cat. 563252), and CD69 BV711 (clone HI.2F3; BD Biosciences, Cat. 740664). Cells were then fixed with intracellular fixation buffer (eBioscience, Cat. 00-8222-49) for 20 min at room temperature, permeabilized (eBioscience, Cat. 00-8333-56), and stained intracellularly for 30 min at 4 °C with the IFN-γ FITC (clone XMG1.2; eBioscience, Cat. 11-7311-82). Fluorescence minus one and isotype controls were included as negative controls. For tetramer staining experiments, H-2D^b^ N_219-227_ tetramer (NIH Tetramer Core) was used.

### ELISAs

Cytokine and chemokine levels in lung homogenates were measured using multiplex ELISA kits on the Meso Scale Discovery (MSD) platform. For samples generated in the BSL-3 facility, lung homogenate supernatants were decontaminated by UV irradiation in accordance with OSU BSL-3 biosafety guidelines.

### Histology and Image Analysis

Lungs were removed from infected mice and fixed in 10% formalin at room temperature. Sample preparation, processing, and hematoxylin and eosin (H&E) staining were performed by Histowiz, Inc. (Brooklyn, NY). H&E images were analyzed for tissue cellularity using a custom R script developed in-house. Briefly, images were converted to grayscale, and global thresholding was applied to identify white spaces. Connected components corresponding to large empty regions were excluded based on a predefined size threshold. Morphological operations (convex hull filling, closing, and dilation) were applied to refine the tissue mask. Adaptive thresholding was then performed within the masked region to identify cellular structures. Cellularity was quantified as the ratio of cellular area to total tissue area, expressed as a percentage.

### RNA extraction, purification and sequencing

Total RNA was extracted from infected lungs by TRIzol reagent (Thermo Fisher Scientific, 15596026) according to the manufacturer’s instructions. RNA cleaning and concentration was done using Zymo Research, RNA Clean & Concentrator-5 kit (cat# R1015) following the manufacturer’s protocol. Fluorometric quantification of RNA and RNA integrity analysis were carried out using RNA Biochip and Qubit RNA Fluorescence Dye (Invitrogen). cDNA libraries were generated using NEBNext Ultra II Directional (stranded) RNA Library Prep Kit for Illumina (NEB #E7760L). Ribosomal RNA was removed using NEBNext rRNA Depletion Kit (human, mouse, rat) (E #E6310X). Libraries were indexed using NEBNext Multiplex Oligos for Illumina Unique Dual Index Primer Pairs (NEB #644OS/L). Library prep generated cDNA was quantified and analyzed using Agilent DNA chip and Qubit DNA dye. Ribo-depleted total transcriptome libraries were sequenced on an Illumina NovaSeq SP flow cell (paired-end 155bp format; 35-40 million clusters, equivalent to 70-80 million reads. Library preparation, QC, and sequencing was carried out at Genewiz.

RNA-seq data were quantified using Salmon, and transcript-level estimates were summarized to gene-level counts with the tximport package. Gene expression analysis was performed with DESeq2 in R, comparing wild-type and Casp11⁻/⁻ samples (n = 4 per group). Low-abundance genes were removed prior to analysis. Differential expression was determined using the Wald test with Benjamini–Hochberg adjustment for multiple testing, and results are reported as log₂ fold change with false discovery rate (FDR). Principal component analysis (PCA) was performed on variance-stabilized data to assess sample clustering. Volcano plots were generated to visualize significantly up- and down-regulated genes, with selected cytokine and chemokine genes highlighted. Expression of key cytokine/chemokine genes was further examined using boxplots of normalized counts. Gene Ontology (GO) enrichment analysis was carried out on significantly up- and down-regulated genes to identify enriched biological processes. Heatmaps of differentially expressed cytokine and chemokine genes were generated to visualize expression patterns across samples.

All analyses and visualizations were performed in R using commonly available Bioconductor and CRAN packages.

### Gene Set Enrichment Analysis (GSEA)

Gene set enrichment analysis was performed with the gseGO function in clusterProfiler, using fold-change values from differential expression without gene filtering. Analyses included all three GO categories (BP, MF, CC) with parameters nPerm = 10,000, minGSSize = 3, maxGSSize = 800, pvalueCutoff = 0.05, and pAdjustMethod = "BH". Normalized enrichment scores (NES) indicate direction and magnitude of enrichment, with NES > 0 denoting activation and NES < 0 denoting suppression. Top 20 enriched terms were visualized by dot plots.

### PCR

Total RNA was extracted and reverse-transcribed to cDNA following the manufacturer’s protocol. Quantitative PCR (qPCR) was performed using SYBR Green chemistry with gene-specific primers for the SARS-CoV-2 nucleocapsid (N) gene, Il1b, Il6, and Cxcl1 (KC). Amplifications were carried out in 10–20 μL reactions on a real-time PCR system under standard cycling conditions. Gapdh was used as the reference gene. Relative copy number was calculated by normalizing target Ct values to Gapdh and then to control samples using the 2^–ΔΔCt method.

### Plaque assay

SARS-CoV-2 plaque assay was performed on Vero E6 TMPRSS2 cells in a 12-well plate incubated for 2 days. Plates were infected with 10-fold serial dilutions of lung. homogenates supernatant. After absorption for 1 h at 37°C, cells were overlaid with 1 ml of DMEM containing 0.25% (w/v) low-melting temperature agarose, 0.12% (v/v) NaHCO3, 2% (v/v) FBS, 25mM HEPES, 2mM L-Glutamine, 100μg/ml of streptomycin, and 100U/ml penicillin. After incubation at 37°C for 24-48 hours cells were fixed with 4% paraformaldehyde for 2 h. The overlay was then removed, and the plaques were visualized by staining with 0.05% (v/v) crystal violet.

### Statistics

Statistical analyses and data visualization were performed using GraphPad Prism Software (version 9.3.1). The specific statistical analyses used are described in the figure legends and include Kaplan-Meier survival analysis, unpaired, 2-tailed t tests, 2-way ANOVA with Šidák’s multiple-multiple comparison test, and 1-way ANOVA with Tukey’s multiple-comparison test. A P value less than 0.05 was considered significant.

## Author contributions

MME, MMS, designed experiments, conducted experiments, analyzed data, and wrote the manuscript. OW, HMA, JA, AB,, JH, JO, GG, YH, RA, RP, SF, ADK, DB, SE, KD, AC, MK, conducted experiments. MP, AW, XZ, analyzed data. MEP, EAH, SMN, ECB, JL, PB, JSY, BS, PD provided critical reagents. AOA conceived of the study, designed experiments, performed experiments, analyzed data, and wrote the manuscript.

## Acknowledgements

We thank the NIH Tetramer Core Facility (contract no. 75N93020D00005) for providing N219 tetramers. This work was supported by NIH grant 5P01AI175399-02 & 5R01HL168501-03.

