## Supplementary material for "Caspase-11 Mediated Hyperinflammation Impairs CD8⁺ T Cell Immunity and Viral Clearance in Severe SARS-CoV-2 Infection": Suppl

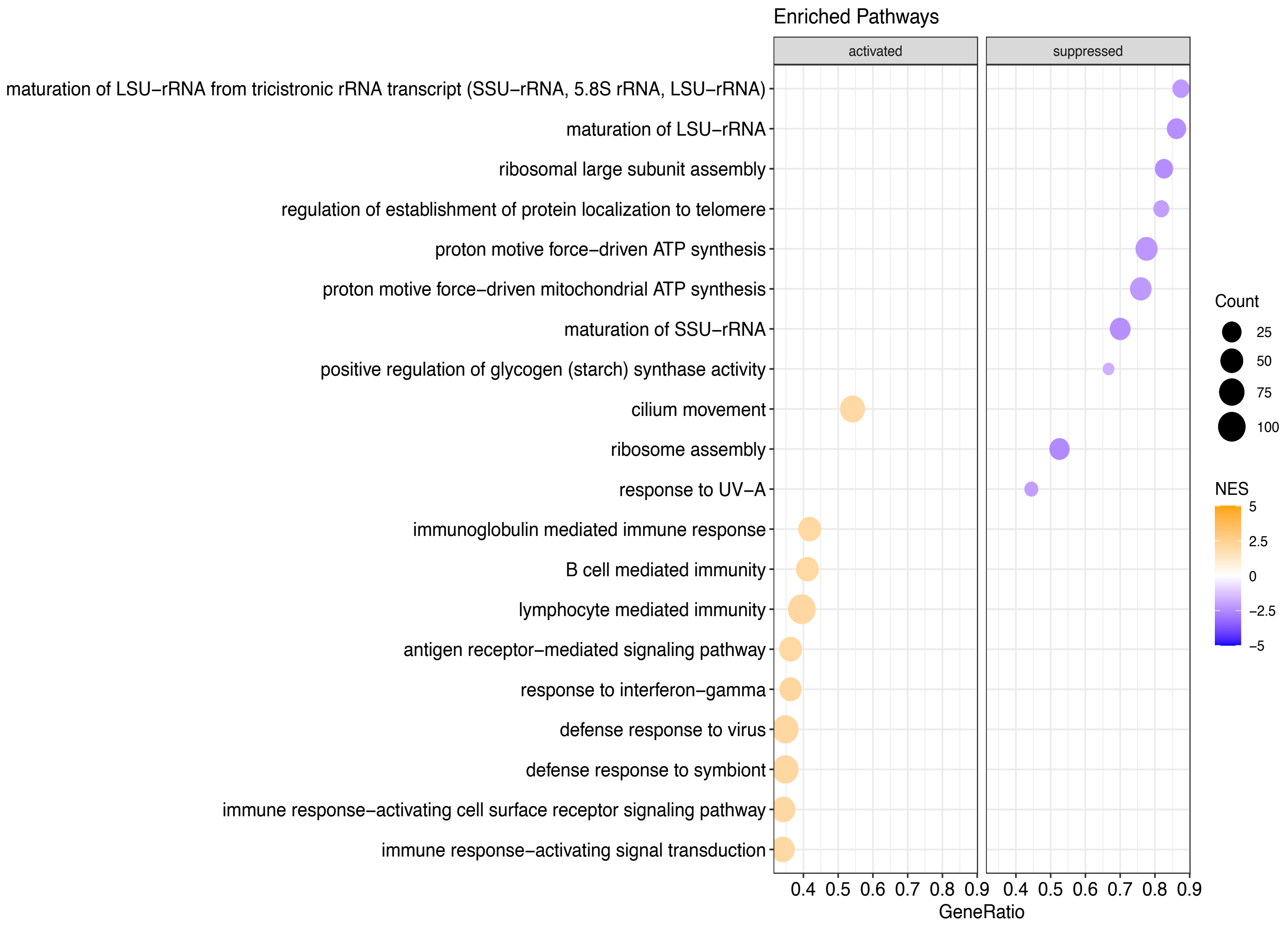

Enriched Pathways
activated
suppressed
maturation of LSU−rRNA from tricistronic rRNA transcript (SSU−rRNA, 5.8S rRNA, LSU−rRNA)
maturation of LSU−rRNA
ribosomal large subunit assembly
regulation of establishment of protein localization to telomere
proton motive force−driven ATP synthesis
proton motive force−driven mitochondrial ATP synthesis
maturation of SSU−rRNA
positive regulation of glycogen (starch) synthase activity
cilium movement
ribosome assembly
response to UV−A
immunoglobulin mediated immune response
B cell mediated immunity
lymphocyte mediated immunity
antigen receptor−mediated signaling pathway
response to interferon−gamma
defense response to virus
defense response to symbiont
immune response−activating cell surface receptor signaling pathway
immune response−activating signal transduction
0.4 0.5 0.6 0.7 0.8 0.9 0.4 0.5 0.6 0.7 0.8 0.9
GeneRatio
Count
25
50
75
100
NES
5
2.5
0
−2.5
−5

**Supplementary Figure 1. CASP11 deficiency enhances antiviral and lymphocyte-mediated immune responses during SARS-CoV-2 infection.**

Gene Set Enrichment Analysis (GSEA) of bulk RNA-seq data from lungs of *Casp11*$^{-/-}$ versus WT mice at day 4 post-infection. Enrichment plots show significant upregulation of pathways related to antiviral immunity, including the interferon-γ response, defense response to virus, and lymphocyte-mediated immunity in *Casp11*$^{-/-}$ group compared to WT group. Additional enriched pathways include B cell- and immunoglobulin-mediated responses, consistent with a shift toward adaptive immune activation in *Casp11*$^{-/-}$ lungs (n = 4/group). Equal number of males and females per group.

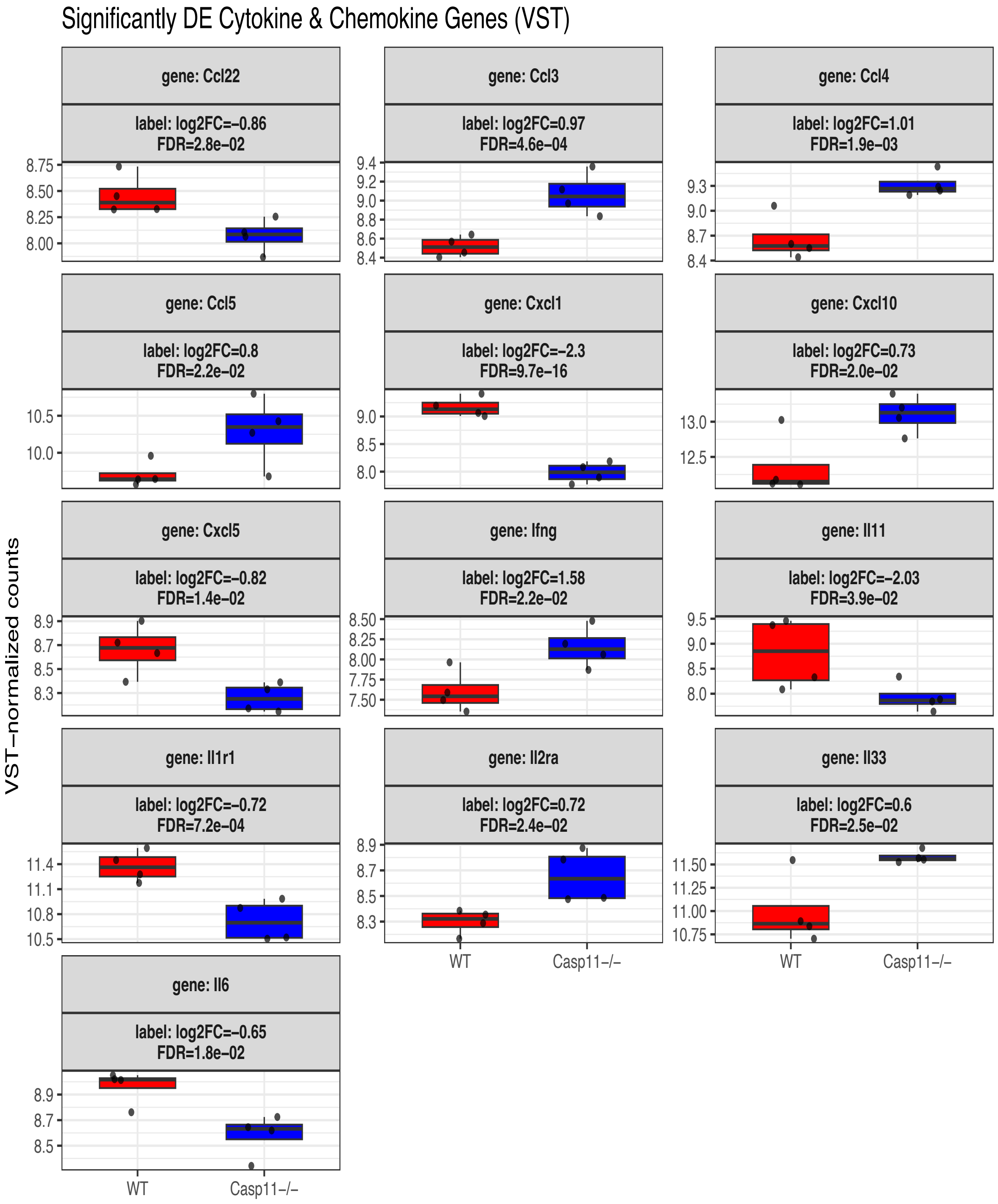


**Supplementary Figure 2. CASP11 deficiency reprograms cytokine and chemokine expression toward lymphocyte-promoting profiles during SARS-CoV-2 infection.**

Expression profiles of significantly differentially expressed cytokine and chemokine genes in lungs of WT and *Casp11*$^{-/-}$ mice at day 4 post-infection. Plots show variance-stabilized transformed (VST) normalized counts and corresponding $\log_2$ fold change values, generated from DESeq2 differential expression (DE) analysis of bulk RNA-seq data using R. *Casp11*$^{-/-}$ lungs exhibited elevated expression of Ifng, Ccl3, Ccl4, Ccl5, and Cxcl10, associated with antiviral responses and effector T cell recruitment. In contrast, genes involved in neutrophil and monocyte recruitment and tissue inflammation, Cxcl1, Cxcl5, Ccl22, Il6, Il11, and Il1r1, were significantly downregulated. (n = 4/group) (Equal number of males and females per group).

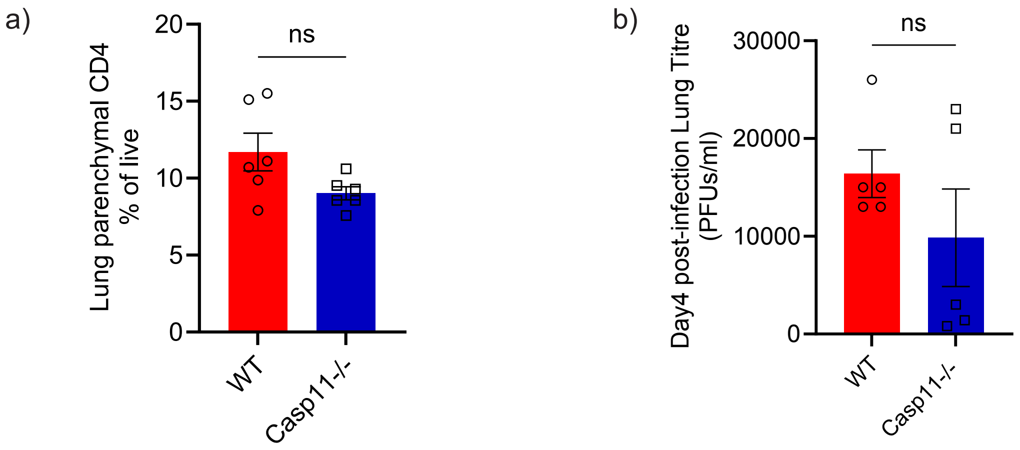


**Supplementary Figure 3. CASP11 has no effect on CD4 T cells or early viral clearance in SARS-CoV-2 infection.**

(a) At day 7 post-infection, flow cytometry with intravascular anti-CD45 labeling revealed no significant difference in lung parenchymal (CD45^lo) CD4$^+$ T cell numbers between *Casp11*$^{-/-}$and WT mice. (n = 6/group) (Equal number of males and females per group). Unpaired t-test. ns, not significant

(b) Viral titers in lung homogenates assessed by plaque assay at day 4 post-infection. Both groups had comparable viral load at day 4 (n = 5/group). ns, not significant

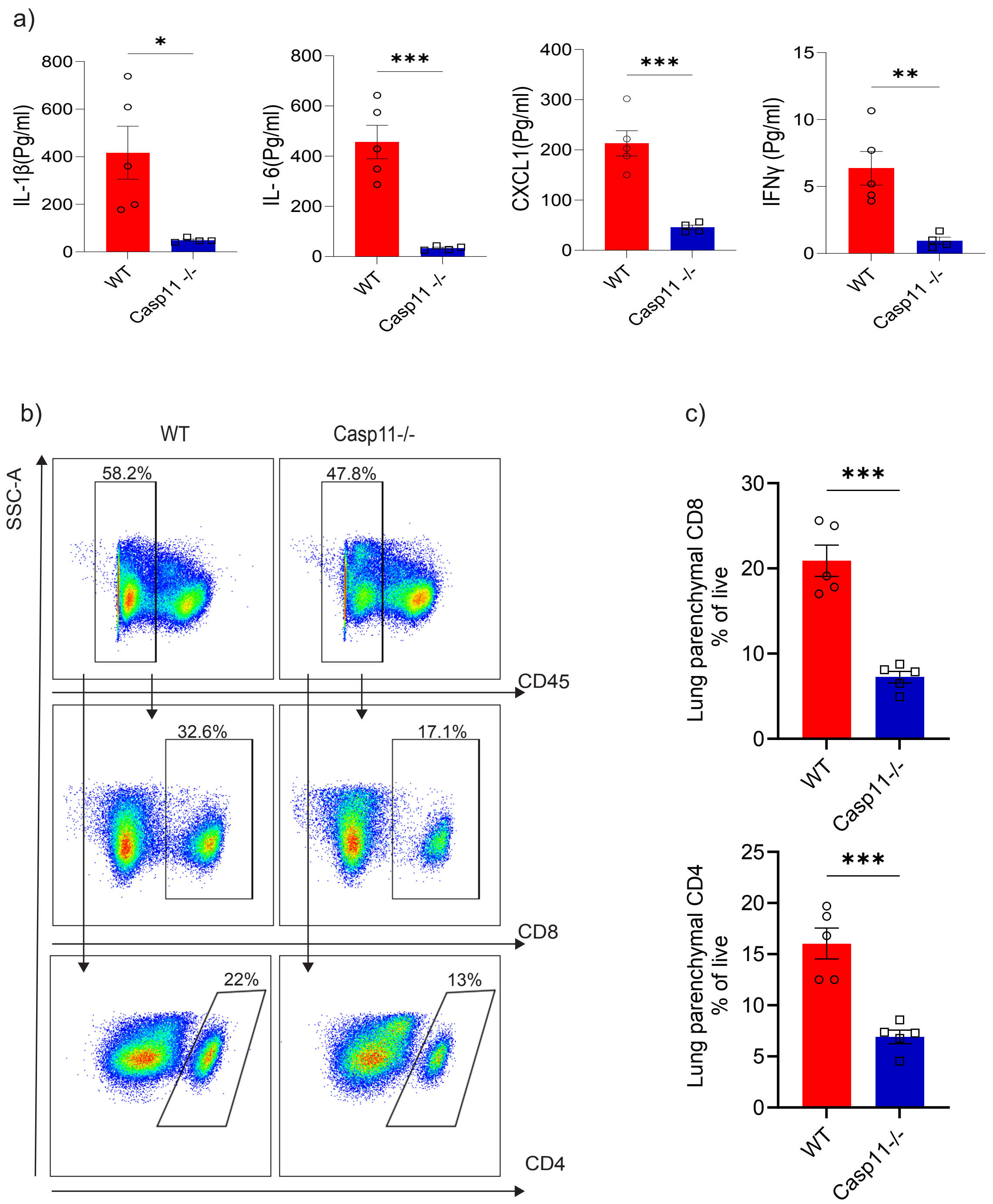
a)
IL-1β(Pg/ml)
*
WT
Casp11 -/-
IL- 6(Pg/ml)
***
CXCL1(Pg/ml)
***
IFNγ (Pg/ml)
**
b)
WT
Casp11-/-
SSC-A
58.2%
47.8%
CD45
32.6%
17.1%
CD8
22%
13%
CD4
c)
Lung parenchymal CD8
% of live
***
Lung parenchymal CD4
% of live
***
Casp11-/-

**Supplementary Figure 4. CASP11 mediates persistent inflammatory and T cell responses after SARS-CoV-2 infection, indicating ongoing inflammation and immune activation despite viral clearance.**

(a) Multiplex ELISA for cytokines at day 14 post infection. (n = 5/ WT, n= 4 *Casp11*$^{-/-}$).

(b–c) Flow cytometric quantification showed increased numbers of lung parenchymal CD8$^+$ and CD4$^+$ T cells in WT compared to *Casp11*$^{-/-}$ mice. Representative flow cytometry plots (b) and quantification (c) are shown. (n = 5/group). unpaired t-test.

*p < 0.05; **p < 0.01; ***p < 0.001

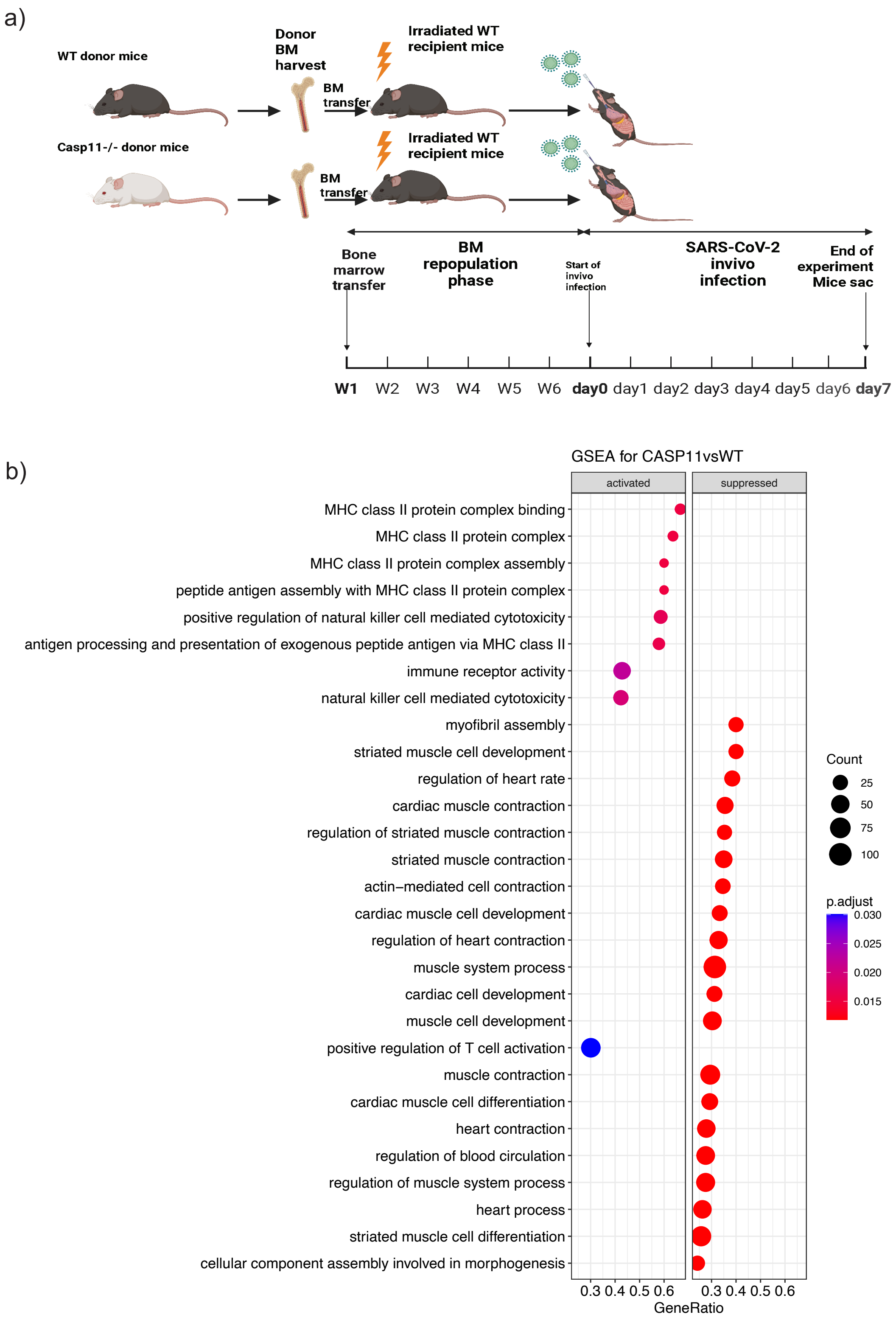
a)
WT donor mice
Casp11-/- donor mice
Donor BM harvest
BM transfer
BM transfer
Irradiated WT recipient mice
Irradiated WT recipient mice
Bone marrow transfer
BM repopulation phase
Start of invivo infection
SARS-CoV-2 invivo infection
End of experiment Mice sac
W1 W2 W3 W4 W5 W6 day0 day1 day2 day3 day4 day5 day6 day7
b)
GSEA for CASP11vsWT
activated
suppressed
MHC class II protein complex binding
MHC class II protein complex
MHC class II protein complex assembly
peptide antigen assembly with MHC class II protein complex
positive regulation of natural killer cell mediated cytotoxicity
antigen processing and presentation of exogenous peptide antigen via MHC class II
immune receptor activity
natural killer cell mediated cytotoxicity
myofibril assembly
striated muscle cell development
regulation of heart rate
cardiac muscle contraction
regulation of striated muscle contraction
striated muscle contraction
actin−mediated cell contraction
cardiac muscle cell development
regulation of heart contraction
muscle system process
cardiac cell development
muscle cell development
positive regulation of T cell activation
muscle contraction
cardiac muscle cell differentiation
heart contraction
regulation of blood circulation
regulation of muscle system process
heart process
striated muscle cell differentiation
cellular component assembly involved in morphogenesis
0.3 0.4 0.5 0.6
0.3 0.4 0.5 0.6
GeneRatio
Count
25
50
75
100
p.adjust
0.030
0.025
0.020
0.015

**Supplementary Figure 5. CASP11 deficiency in hematopoietic cells enhances survival, reduces inflammation, and promotes T cell responses after SARS-CoV-2 infection.**

(a) Schematic of bone marrow (BM) chimera generation. Lethally irradiated WT recipients were reconstituted with either WT or *Casp11*$^{-/-}$donor bone marrow, creating WT→WT and *Casp11*$^{-/-}$→WT (*Casp11*$^{-/-}$ chimera) mice. This setup enabled assessment of hematopoietic CASP11 function while maintaining WT non-hematopoietic compartments, with alveolar macrophages remaining WT due to their radioresistant, self-renewing nature.

(b) Gene set enrichment analysis (GSEA) of RNA-seq data from lungs harvested at day 4 post-infection revealed upregulation of T cell activation and antigen presentation pathways in *Casp11*$^{-/-}$ chimeras, along with suppression of pathways related to muscle and contraction processes. (n = 4/group) (Equal number of males and females per group).

a)

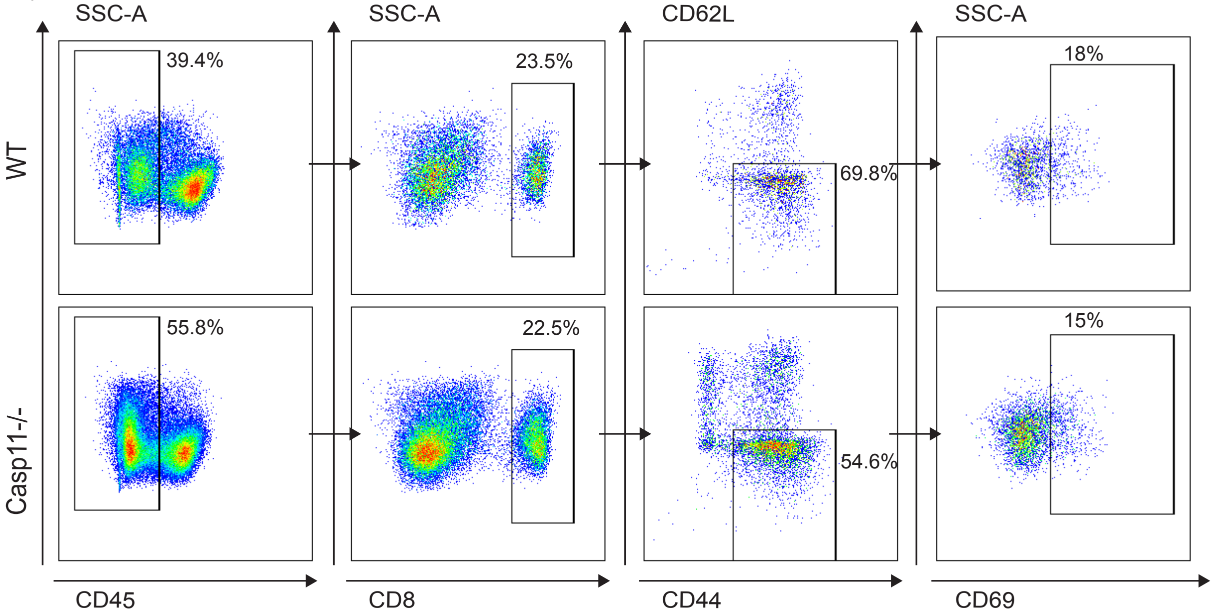
SSC-A
SSC-A
CD62L
SSC-A
WT
Casp11-/-
39.4%
55.8%
23.5%
22.5%
69.8%
54.6%
18%
15%
CD45
CD8
CD44
CD69

b)

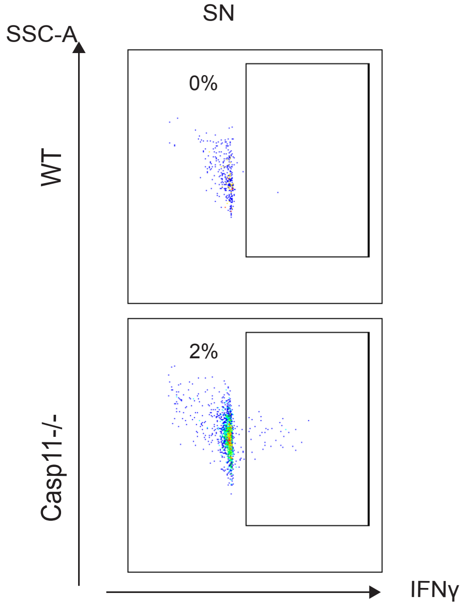
SSC-A
SN
WT
Casp11-/-
0%
2%
IFNγ

**Supplementary Figure 6. Gating strategy of flow cytometry analysis of CD8 T cells in infected lungs of Chimera mice.**

(a) Flow cytometric analysis of lung parenchymal CD8$^+$ T cells and activated effector CD8$^+$ T cells (CD62L^loCD44^hiCD69^hi) at day 7 post-infection.

(b) IFN-γ$^+$ CD8$^+$ T cells following ex vivo stimulation with spike+nucleocapsid (SN) peptide mix**.**

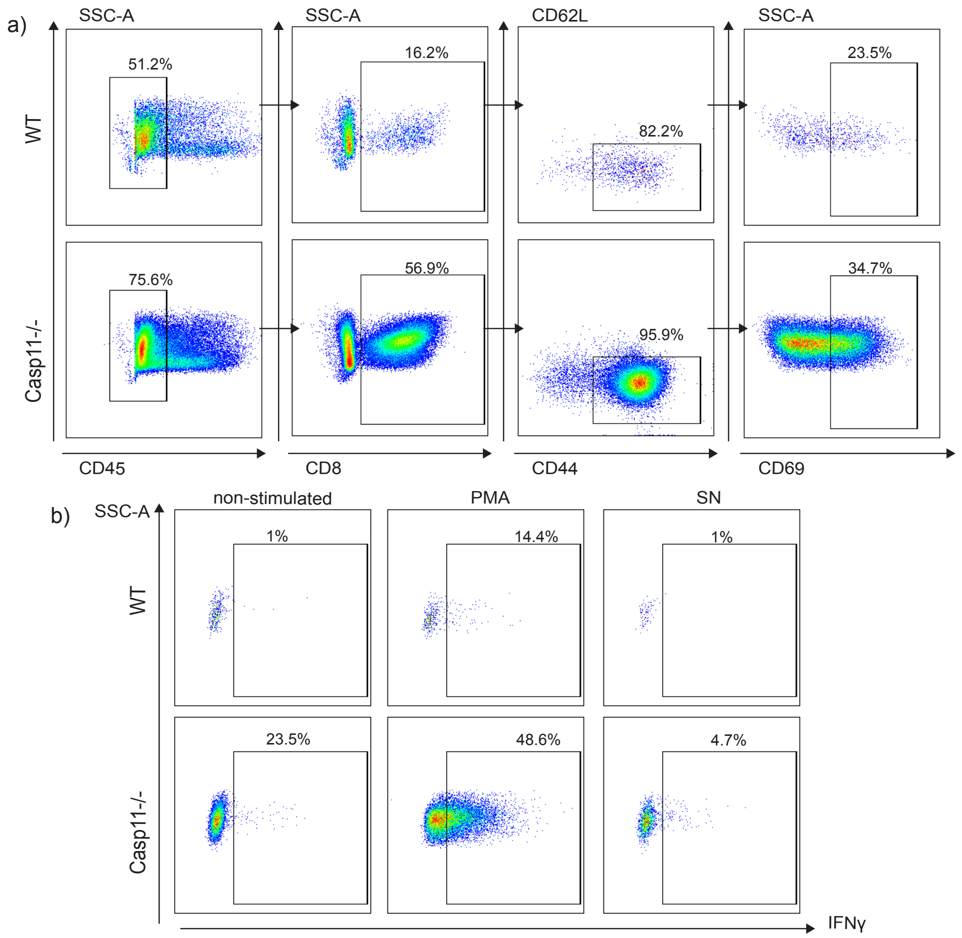


**Supplementary Figure 7. Gating strategy of flow cytometry analysis of CD8 T cells in infected lungs of *Casp11*$^{-/-}$ flox mice.**

(a) Flow cytometric analysis of lung parenchymal CD8$^+$ T cells and activated effector CD8$^+$ T cells (CD62L^loCD44^hiCD69^hi) at day 7 post-infection

(b) IFN-$\gamma^+$ $CD8^+$ T cells following ex vivo stimulation with PMA/Ionomycin or spike+nucleocapsid (SN) peptide mix.
